# Microbiome shifts associated with the introduction of wild Atlantic horseshoe crabs (*Limulus polyphemus*) into a touch-tank exhibit

**DOI:** 10.1101/2020.02.27.968784

**Authors:** Ariel D. Friel, Sean A. Neiswenter, Cale O. Seymour, Lauren Rose Bali, Ginger McNamara, Fabian Leija, Jack Jewell, Brian P. Hedlund

## Abstract

The Atlantic horseshoe crab (*Limulus polyphemus*) is a common marine aquarium species and model organism for research. There is potential monetary and conservation value in developing a stable captive population of horseshoe crabs, however, one major impediment to achieving captivity is a lack of knowledge regarding captive diseases. We utilized 16S rRNA gene amplicon sequencing to track changes in the microbiomes of four body locations in three wild-caught (tracked over 14 months in captivity) and three tank-acclimated (>2 years in captivity) adult *L. polyphemus* in a touch tank at Shark Reef Aquarium at Mandalay Bay in Las Vegas, NV. The wild population hosted diverse and distinct microbiomes on the carapace (737 amplicon sequence variants or ASVs), cloaca (1078 ASVs), gills (941 ASVs), and oral cavity (1193 ASVs), which were dominated by classes *Gammaproteobacteria*, *Bacteroidia*, and *Alphaproteobacteria*. A rapid decline in richness across all body locations was observed within 1 month of captivity, with tank-acclimated (>2 years) animals having <5% of the initial microbiome richness and a nearly completely restructured microbial community. The carapace, oral cavity, and gills of the tank-acclimated animals hosted abundant populations of *Aeromonas* (>60%) and *Pseudomonas* (>20%), both of which are known opportunistic pathogens of aquatic animals and can express chitinases, providing a plausible mechanism for the development of the carapace lesion pathology observed in this and other studies. These results provide an important baseline on the microbiomes of both wild and tank-acclimated horseshoe crabs and underscore the need to continue to investigate how native microbial populations may protect animals from pathogens.

## Introduction

Perhaps the most challenging environmental change an organism can experience is when a wild individual is taken from a natural setting and held in captivity under artificial conditions as is the case in a laboratory, the pet trade, the food industry, or zoos and aquaria. Replicating the natural environment is nearly impossible under artificial conditions and depending on the circumstances it may be necessary or more convenient to modify conditions, such as temperature or salinity, from those an organism would experience in the wild. Additionally, although some conditions might be controlled to limit variability under captive conditions, other variables might cycle or build up to unnatural levels in captivity, such as the case with dissolved nitrogen in aquaculture (Hargreaves 1998). In captivity an organism may also be exposed to population densities and different species they would never encounter in natural settings, fostering novel biotic interactions. This can be particularly pronounced for aquatic organisms, such as in aquaculture or large aquaria, where high densities of a variety of host species share the same tank or have a common source of filtered water. In such artificial systems stress can lead to microbiome dysbiosis and infections by obligate or opportunistic pathogens (Llewellyn et al. 2014).

*Limulus polyphemus*, the Atlantic horseshoe crab, is one of four species in the order *Xiphosurida* and the only species in North America. Horseshoe crabs have a deep evolutionary history dating over 400 mya to at least the Ordovician (Rudkin et al 2008). Contrary to what their common name implies, horseshoe crabs are not crustaceans, but represent a highly divergent lineage that is more closely related to sea spiders and other arachnids than crabs (Ballesteros and Sharma 2019). *L. polyphemus* is widespread along the continental shelf of North America’s east coast and is typically regarded as an ecological generalist, in that it occupies a broad range of temperature and salinity. Adults are strictly benthic and burrow through sediments to feed on polychaetes, bivalves, and other benthic fauna. Given their deep evolutionarily history and highly variable ecology it is likely that horseshoe crabs harbor a unique and diverse microbiome. However, to date, only a single study has described microorganisms from wild horseshoe crabs. In that study, bacteria were isolated from the gills and mouth from two species of Asian horseshoe crabs, *Tachypleus gigas* and *Carcinoscorpius rotundicauda*, and identified as members of the genera *Pseudoalteromonas*, *Vibrio*, and *Photobacterium* (Ismail et al., 2015).

*Limulus polyphemus* have considerable value both commercially and ecologically. The pharmaceutical industry utilizes their blood to produce *Limulus* amebocyte lysate, which is used to detect endotoxins and for bait by the commercial eel industry. *L. polyphemus* can also serve as a model organism for a variety of research topics, including embryology and vision research (Liu and Passaglia 2009; Williams 2019). Additionally, millions of migratory birds refuel on their eggs each spring at spawning sites (Smith et al., 2006). As a highly unusual and non-threatening organism, they are also used to garner interest and educate the general public through interactive exhibits at aquaria.

Due to multifaceted threats, horseshoe crab populations have been in decline (Smith et al. 2017). There is a potential utility to multiple stakeholders in the maintenance of captive populations (Carmichael and Brush 2012). However, very little is known of the microbial communities of either healthy or diseased horseshoe crabs. A common affliction of captive horseshoe crabs are lesions that occur on the carapace (Nolan and Smith 2009). These lesions have been associated with a variety of microorganisms including heterotrophic bacteria (e.g., Thompson et al., 2011), *Cyanobacteria* (Leibovitz 1986), fungi (Tuxbury et al., 2014), and green algae (Braverman et al., 2012). However, no systematic studies have explored the microbiomes of wild or captive horseshoe crabs using cultivation-independent techniques.

The main objective of this study was to document the microbiomes of wild, adult horseshoe crabs at several body locations and document the progression of microbial dysbiosis following their captivity. We further extended the study by examining the same body locations on several long-term captive animals (>2 years) in the same tank. From this, we hoped to identify possible symbionts or commensals of a wild horseshoe crab microflora as well as potential pathogens that develop *ex situ*. Our study revealed highly diverse microbial communities in the carapace, cloaca, gills, and oral cavity of wild animals and documented a rapid and steep decline in microbial richness and near-complete restructuring of the microbial community following captivation. The opportunistic pathogens *Aeromonas* (>60%) and *Pseudomonas* (>20%) together comprised >80% of the microbiomes of animals acclimated to the aquarium for over two years. In contrast, the cloaca of the tank-acclimated animals was distinct and more diverse, with abundant populations of *Aeromonas*, *Shewanella*, *Vagococcus*, and *Enterococcus*. This study forms a baseline for both wild and captive horseshoe crabs and provides a timeline and body atlas for development of dysbiosis. Potential mechanisms for maintenance of wild and diverse microbiota, and their potential importance in health, are discussed herein.

## Materials and Methods

### Sampling and Experimental Design

Three wild animals and native sediments were collected from the wild and dry-shipped overnight by Dynasty Marine (Marathon, Florida). Upon receipt, the carapace, book gills, oral cavity, and cloaca were sampled with sterile swabs. The naïve individuals were then uniquely identified by attaching a PIT tag to the underside of their carapace with epoxy and introduced into the public touch-tank exhibit at the Shark Reef Aquarium at Mandalay Bay Resort and Casino in Las Vegas, Nevada. At one month, nine months, and 14 months following introduction to the touch tank, the same animals were sampled at the same four locations using the same protocol as described above. One wild-caught cloacal sample was not collected at the one-month time point. Prior to introducing the naïve individuals into the touch tank, the substrate at the bottom of the tank and three tank-acclimated horseshoe crabs (> 2 years in captivity) were sampled at the same body locations and using the same protocol to establish a baseline of the microbial community already present. Swabs were immediately taken to the lab for DNA isolation, described below. Due to a laboratory error during DNA isolation one tank-acclimated horseshoe crab’s gill sample was not further processed.

### Captive Conditions

Captive horseshoe crabs were maintained in the public touch-tank exhibit at the Shark Reef Aquarium at Mandalay Bay Resort and Casino (Las Vegas, NV) during the study. The tank is 2,500 gallons with a water depth of 34 cm and 4 cm of fine aragonite sand for substrate. The water is held at 23ºC, pH 8.1, and a salinity of 33 ppt using Instant Ocean (Blacksburg, VA). The light cycle is 16 hours of dim indirect ambient lighting similar to dusk conditions and 8 hours of dark. Filtration includes pressure sand-filters, foam fractionator, and a trickle de-gassing bio-filter with twice weekly filter backwashes and 10% water changes. Horseshoe crabs are fed a combination of earthworms, clams, Superba krill, and oysters six days per week. Any uneaten food is immediately removed from the tank. Up to 9 individual horseshoe crabs occupied the tank during the present study. Other species that share the tank include coral catsharks, *Atelomycterus marmoratus,* and several genera of rays such as *Neotrygon*, *Glaucostegus*, *Urobatis*, and *Trygonorrhina*.

### 16S rRNA gene amplicon sequencing

DNA was extracted from horseshoe crab samples taken using sterile swabs and substrate samples (all stored at −80 °C) using the FastDNA™ SPIN Kit for Soil (MP Biomedicals, Santa Ana, CA) according to manufacturer’s instructions. The V4 region of the 16S rRNA gene was amplified and sequenced using the updated Earth Microbiome Project bacterial- and archaeal-specific 515F/806R primer set (Thompson et al., 2017). Amplification, library preparation, and sequencing were performed at Argonne National Laboratory (Lemont, IL) on an Illumina MiSeq platform (2×151 bp).

### Sequence processing, quality control, and data

Paired-end Illumina MiSeq reads were quality filtered, aligned, and assigned to amplicon sequence variants (ASVs) using DADA2 (Callahan et al., 2016) via Qiime2 version 2019.1 (Boylen et al., 2018, Caparaso et al., 2010). ASVs were classified in Qiime2 using a naïve-Bayesian classifier (Bokulich et al. 2018) trained on the V4 region of the Silva NR99 132 alignment (Pruesse et al 2007). Those classified as chloroplast or mitochondrial were removed.

### Microbial community data analysis

ASVs and Silva-based taxonomy were exported from Qiime2 to be analyzed using R. To normalize the samples for highly variable sequencing depth, ASV counts were scaled to counts-per-million. Diversity indices (Faith’s PD, Observed, Shannon, and InvSimpson) were calculated using R packages phyloseq version 1.28.0 (McMurdie and Holmes 2013) and picante version 1.8 (Kimbel et al. 2010). The phylogenetic tree used to calculate UniFrac and Faith’s PD diversity metrics was generated using FastTree (Price et al. 2009, 2010) in Qiime2 with the developer-recommended settings. Between-sample dissimilarity was calculated using the Bray-Curtis algorithm implemented in R package vegan version 2.5-6 (Dixon 2003). Ordination was performed using non-metric multidimensional scaling (NMDS) via the R packages phyloseq and vegan. ASVs were agglomerated at the family-level using the tax_glom function from phyloseq and regressed against each distance matrix using the envfit function of vegan. Taxonomic vectors representing the bacterial and archaeal families that were significantly (p-value < 0.05) correlated with community dissimilarity between samples, as determined by envfit, were displayed on the NMDS plot. To analyze the similarity in ASV composition over time in captivity, the community matrix was combined by body location to generated Venn diagram bins. To further understand which specific ASVs were changing over time in captivity, differential abundance analysis was conducted using DESeq2 version 1.24.0 (Love, Huber, and Anders 2014). Differentially abundant ASVs of p < 0.01 and p < 0.05 were aligned using the SINA alignment tool (Pruesse et al. 2012). A phylogenetic tree was constructed from these using IQ-Tree 1.6.7.a (Minh et al. 2013, Nguyen et al. 2015, Kalyaanamoorthy 2017), rooted at its midpoint phytools 0.6.60 (Revell 2011), and ladderized using ape 5.3 (Paradis et al. 2004). All figures were rendered using Microsoft PowerPoint, the R package vegan version 2.5-6 (Dixon 2003), or the R package ggplot2 version 3.2.1 (Wickham 2011).

## Results

### Rarefaction Curves and Community α-Diversity

Following quality filtering, 1,758,652 total DNA sequences were recovered from 61 samples (59 horseshoe crab samples and two substrate samples). Of these >1.7 million high-quality DNA sequences, 6,844 unique amplicon sequence variants (ASVs) were identified, comprising 64 bacterial and archaeal phyla. Rarefaction curves for all samples plateaued at a reasonable sequence depth (~5,000 sequences), indicating adequate coverage of the total diversity (Supplemental Figure 1).

**Figure 1.**
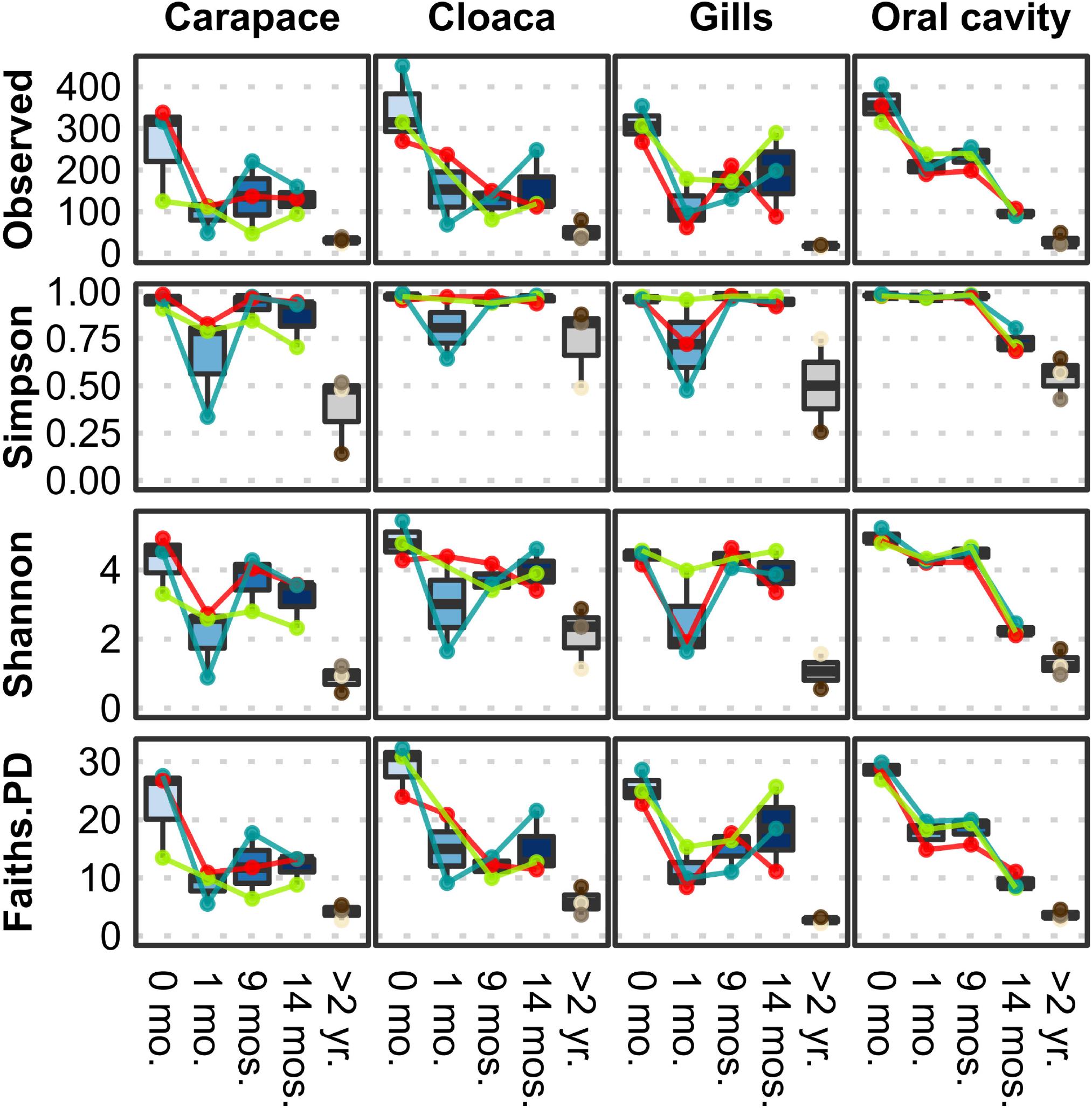
Microbial alpha diversity measures at different locations from wild-caught *Limulus polyphemus* (0 months) introduced to a captive environment and sampled after 1, 9, and 14 months of captivity. Tank-acclimated (>2 years) individuals were sampled prior to introducing the wild-caught individuals into the tank to establish the microbiome already present in the tank. Each colored line represents one of the three wild individuals that was followed over time.

To evaluate the effect of captivity on their microbiomes, several α-diversity indices were calculated at 1 month, 9 months, and 14 months in captivity (Figure 1). At all body locations, the wild horseshoe crabs hosted highly diverse microbial communities, with the mean observed ASVs exceeding 300 and Simpson’s evenness above 0.98. After one month in captivity, a decline was observed for all diversity indices, indicating a loss of richness (observed ASVs), diversity (Simpson and Shannon indices), and phylogenetic diversity (Faith’s PD). The decline was most clear in the observed ASVs and Faith’s PD index at all body locations and richness did not recover through the end of the study, except for partial recovery in the gills of two animals. A similar pattern of severe decline in Simpson’s Evenness was observed after one month for the carapace and gill samples; however, these values returned to near-normal levels within nine months and remained high through the end of the study. The oral cavity samples displayed a progressive loss of richness over the course of the experimental timeline; evenness remained high but started to decline 14 months after captivity. Throughout the 14 months of captivity, the richness and evenness of the wild horseshoe crab microbiomes never reached the drastically low values of the tank-acclimated population that was in captivity for over two years, which had <50 ASVs at each body location.

### Community Composition of Horseshoe Crabs and Substrate

Both wild and tank-acclimated horseshoe crab microbiomes were dominated by *Bacteria* (>99% of ASVs), whereas *Archaea* were more abundant (~3-6%) in the Florida and Shark Reef touch tank substrate samples (Supplemental Figure 2). The Florida substrate was the most diverse of all the samples and was comprised of *Proteobacteria*, *Bacteroidetes*, *Chloroflexi*, *Planctomycetes*, *Spirochaetes*, *Cyanobacteria*, *Calditrichaeota*, *Acidobacteria*, and *Actinobacteria* (Supplemental Figure 3). In comparison, the Shark Reef substrate sample was primarily comprised of *Proteobacteria*, *Bacteroidetes*, *Actinobacteria*, *Nitrospirae*, *Thaumarchaeota*, *Acidobacteria*, and *Planctomycetes*. The phylum- and class-level composition of the wild horseshoe crab microbiomes at the four body locations was highly similar, being dominated by *Proteobacteria* (*Gammaproteobacteria* and *Alphaproteobacteria*) and *Bacteroidetes* (*Bacteroidia*), with individual horseshoe crabs possessing varying abundances of *Planctomycetes*, *Patescibacteria*, *Verrucomicrobia*, and *Actinobacteria*. The phylum- and class-level composition of the wild horseshoe crab microbiome was generally retained throughout captivity, with *Gammaproteobacteria, Bacteroidia*, and *Alphaproteobacteria* being the most abundant classes across nearly all body locations and sampling times. However, over the course of 14 weeks in captivity, transient blooms of *Firmicutes*, *Tenericutes*, *Epsilonbacteraeota*, *Fusobacteria*, and an unclassified bacterial phylum were observed, during which these phyla comprised 8.0% to >70% of the microbiome in individual samples. The majority of the tank-acclimated horseshoe crab samples (9/12) were dominated by *Gammaproteobacteria* (>90% of the total community composition). The other three, all cloaca samples, also contained abundant *Firmicutes*, *Bacteroidetes*, and *Epsilonbacteraeota*. Bar plots showing the relative abundance at the domain, phylum, class, order, family, and genus levels for all samples can be found in the supplemental material (Supplemental Figures 2–7).

**Figure 2.**
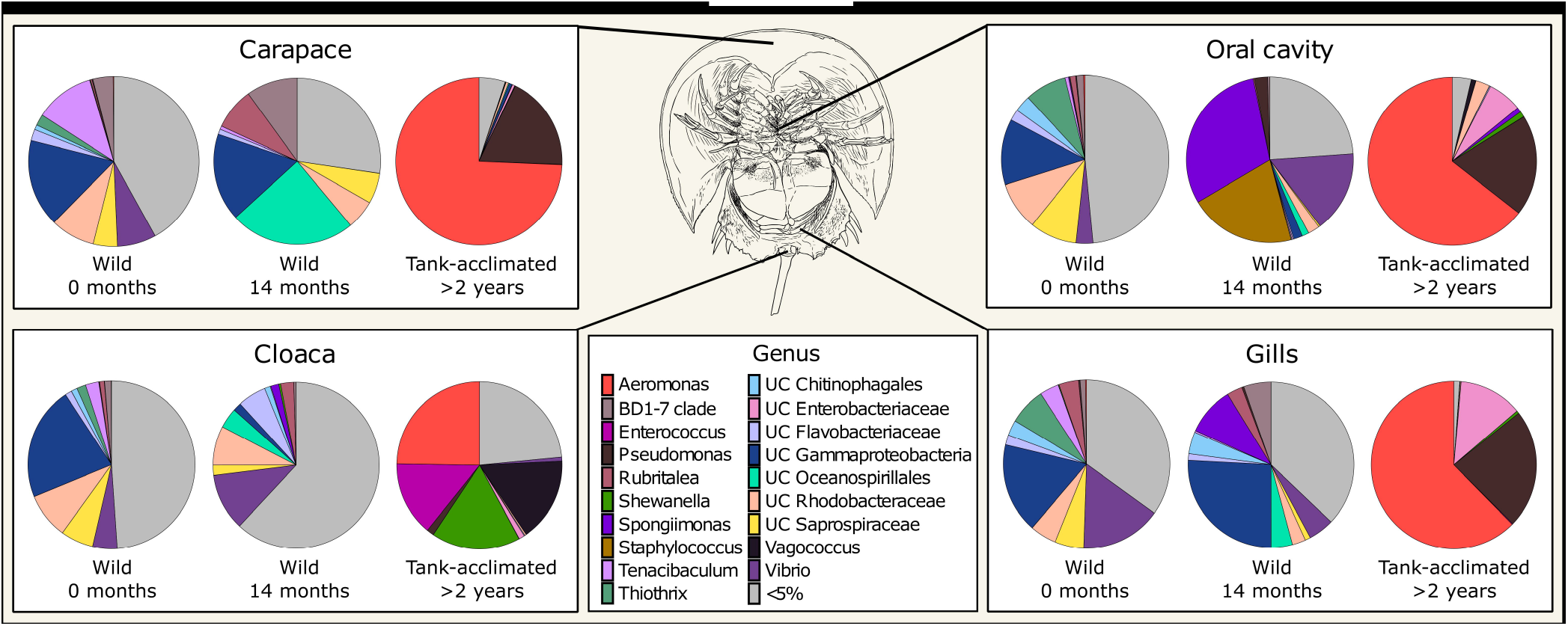
Mean normalized abundance (n=3) of the most abundant genera from different body locations of *Limulus polyphemus*. Wild-caught animals were sampled in the field (0 months) and following 14 months in captivity. For simplicity, samples taken at one and nine months are not shown. The tank-acclimated individuals had been in the tank for over two years but were sampled at the same time the wild-caught 0-month samples were taken to establish the microbiome already present in the tank. Colors represent genera representing >5% of the total microbiome, while gray represents all the uncommon taxa (each <5%) combined.

**Figure 3.**
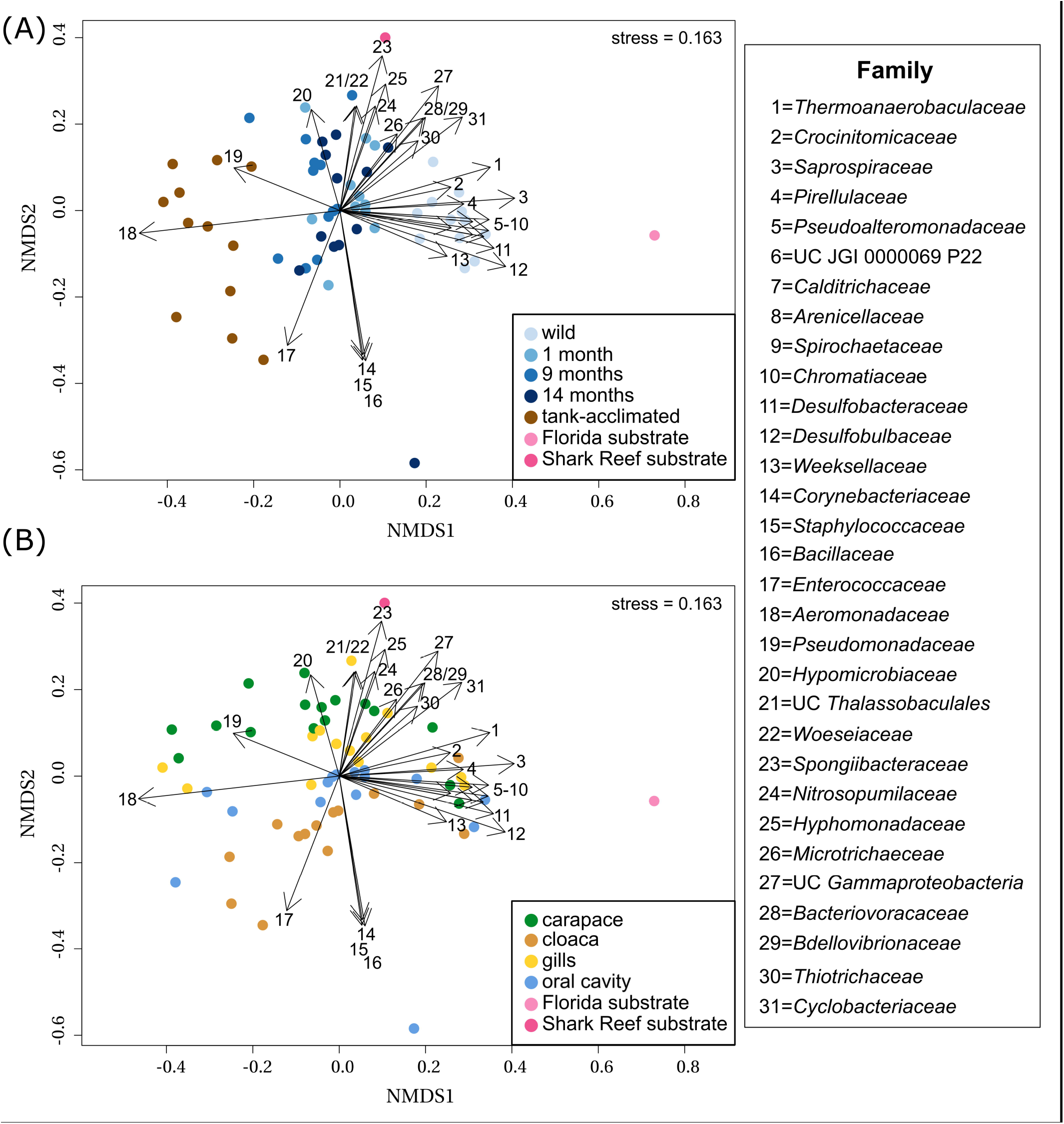
(A) NMDS showing the separation of microbiomes from combined sampling locations over time in captivity along NMDS1, including three wild-caught individuals at different sampling times (blue shades) and three tank-acclimated animals (brown; >2 years in captivity). For reference, substrates from the captive habitat and representative sediments from where the wild caught individuals were captured are shown (pink shades). (B) NMDS showing the separation of microbiomes from combined sampling times by body location along NMDS2. Taxonomic vectors representing bacterial and archaeal families that were significantly (p-value = 0.05) correlated with community dissimilarity were overlaid on to both plots.

Describing phylum-level composition is useful in creating a broad picture of a community and is common practice in the literature, however, it is difficult to extract meaningful information about ecological functions at that taxonomic level. To address this, we explored lower taxonomic levels (Figure 2 and Supplemental Figure 7). The four body locations sampled on the wild horseshoe crabs had highly similar microbial communities, with the most abundant members (at least 50% of the total community) being unclassified members of the *Gammaproteobacteria*, *Chitinophagales*, *Rhodobacteraceae*, *Saprospiraceae*, *Flavobacteriaceae,* and BD1-7 clade, along with the genera *Vibrio*, *Tenacibaculum*, *Thiothrix*, and *Rubritalea* (Figure 2).

Although some dominant genera were retained in the wild-caught population after 14 months in captivity, there were large shifts in community composition, and a divergence of microbial community structure by body location (Figure 2). All body locations were colonized by unclassified *Oceanospirillales* and had a complete loss of *Thiothrix* and decreased abundance of *Tenacibaculum* and unclassified *Rhodobacteraceae* through time. The carapace and cloaca microbial communities had an increased abundance of the genus *Rubritalea*. *Spongiimonas* was observed in the oral cavity, cloaca, and gills, but not the carapace. The composition of the oral cavity samples was drastically different than the initial sample, with an increased abundance of *Vibrio* and an appearance of *Staphylococcus*. Cloaca samples had an increased abundance of *Vibrio* and unclassified *Flavobacteriaceae* and a decreased abundance of unclassified *Gammaproteobacteria*. Gill samples showed an increased abundance of unclassified *Gammaproteobacteria*, *Flavobacteriaceae*, and BD1-7 clade and a decrease in the abundance of *Vibrio*. Over the course of the experimental timeline, transient bloom events by *Shewanella*, *Chromohalobacter*, *Pseudomonas*, *Cocleimonas*, *Bdellovibrio*, and *Prevotella* 9 were observed (Supplemental Figure 6).

*Aeromonas* and *Pseudomonas* dominated the microbial communities of tank-acclimated carapace, oral cavity, and gills, comprising more than 75% of the total community (Figure 2). Additionally, *Cocleimonas* was observed on the carapace and unclassified *Enterobacteriaceae* in the oral cavity and gills (Supplemental Figure 6). The cloaca microbial communities were distinct and more diverse than the other body locations, with *Aeromonas*, *Shewanella*, *Vagococcus*, *Enterococcus*, *Lactococcus*, and *Proteus* present.

### Community Dissimilarity Analysis

A NMDS analysis of Bray-Curtis dissimilarity for all samples enabled visualization of differences in microbial community composition by time and body location (Figure 3). The distinct microbial communities observed across time and body location were significantly different via PERMANOVA (p-value = 0.001 for both); individual was not a significant factor (p-value = 0.124). Samples separated by time along the x-axis (NMDS1) and body location along the y-axis (NMDS2). There was an increasing severity of dysbiosis afflicting the horseshoe crabs following time in captivity, shown by movement from right to left in the NMDS plot (Figure 3A). Both substrate samples were distinct from all horseshoe crab samples and each other, yet the wild horseshoe crab and captive horseshoe crab microbiomes (1-14 months) were most closely related to the Florida and aquarium substrates, respectively, suggesting environmental filtering. The dysbiotic tank-acclimated animal microbiomes were distinct and distant from both substrate microbial communities. As noted previously, the community composition of wild horseshoe crabs was highly similar across all body locations, and it became increasingly structured by body location during captivity, as evidenced by increased distance between points from right to left in the NMDS (Figure 3B). In striking contrast to the initial samples from the wild-caught animals, the community composition of the tank-acclimated horseshoe crabs was highly dissimilar between individuals, yet still retained some structuring by body location.

To gain a deeper understanding of the taxonomic changes associated with captivity and body location, taxonomic vectors representing the bacterial and archaeal families that were significantly (p-value ≤ 0.05) correlated with community dissimilarity between the samples were fitted onto the NMDS ordination (Figure 3). Unique marker taxa were associated with the different horseshoe crab populations and sediment samples. *Pirellulaceae* (#4)*, Pseudoalteromonadaceae* (#5), unclassified JGI 0000069P22 (#6), *Desulfobacteraceae* (#11), *Desulfobulbaceae* (#12), *Weeksellaceae* (#13), *Nitrosopumilaceae* (#24), and *Thiotrichaceae* (#30) were all present in the wild horseshoe crabs but were either partially or completely lost during captivity. Several families were observed at a low abundance in the wild horseshoe crabs but increased significantly during captivity, including *Hyphomicrobiaceae* (#20), *Spongiibacteraceae* (#23), *Hyphomonadaceae* (#25), *Microtrichaeceae* (#26), unclassified *Gammaproteobacteria* (#27), *Bacteriovoracaceae* (#28), and *Bdellovibrionaceae* (#29). The oral cavity of several wild-caught animals after 14 months of captivity hosted blooms of *Corynebacteriaceae* (#14), *Staphylococcaceae* (#15), and *Bacillaceae* (#16). The tank-acclimated animals were dominated by *Enterococcaceae* (#17), *Aeromonadaceae* (#18), and *Pseudomonadaceae* (#19). Several families were associated with the native Florida substrate including *Thermoanaerobaculaceae* (#1), *Calditrichaeceae* (#7), *Spirochaetaceae* (#9), *Chromatiaceae* (#10), *Desulfobacteraceae* (#11), *Desulfobulbaceae* (#12), and *Cyclobacteriaceae* (#31). The touch tank substrate included unclassified *Thalassobaculales* (#21), *Woeseiaceae* (#22), *Spongiibacteraceae* (#23), *Nitrosopumilaceae* (#24), *Hyphomonadaceae* (#25), *Microtrichaeceae* (#26), and *Cyclobacteriaceae* (#31).

### Differential Abundance Analysis of Sequence Variants

To more deeply resolve the shifts occurring in the microbial community structure wild horseshoe crabs as they transitioned to captivity, we conducted several ASV-focused analyses. First, the number of ASVs exclusive to and shared between timepoints was compared (Figure 4). The ASV composition of each time grouping was largely unique (>80%), with very little overlap between time points or between the wild and tank-acclimated populations. Very few ASVs were present throughout the study (0.2-0.6%), underscoring the dynamic nature of the microbiome during the transition to captivity. Thus, even though the broader taxonomic structure (i.e. phylum and class) of the wild horseshoe crab microbiome was retained throughout captivity, the microbiome was nearly completely restructured at the ASV level. The loss of diversity evident in the alpha diversity measurements (Figure 1) was also evident here, as evidenced by the large proportion of unique ASVs in the wild-caught population (44%-51% of the total ASV composition), and the decline of unique ASVs over time in captivity. The tank-acclimated horseshoe crab population had the lowest number of unique ASVs (0.8%-5.5% of the total ASV composition).

**Figure 4.**
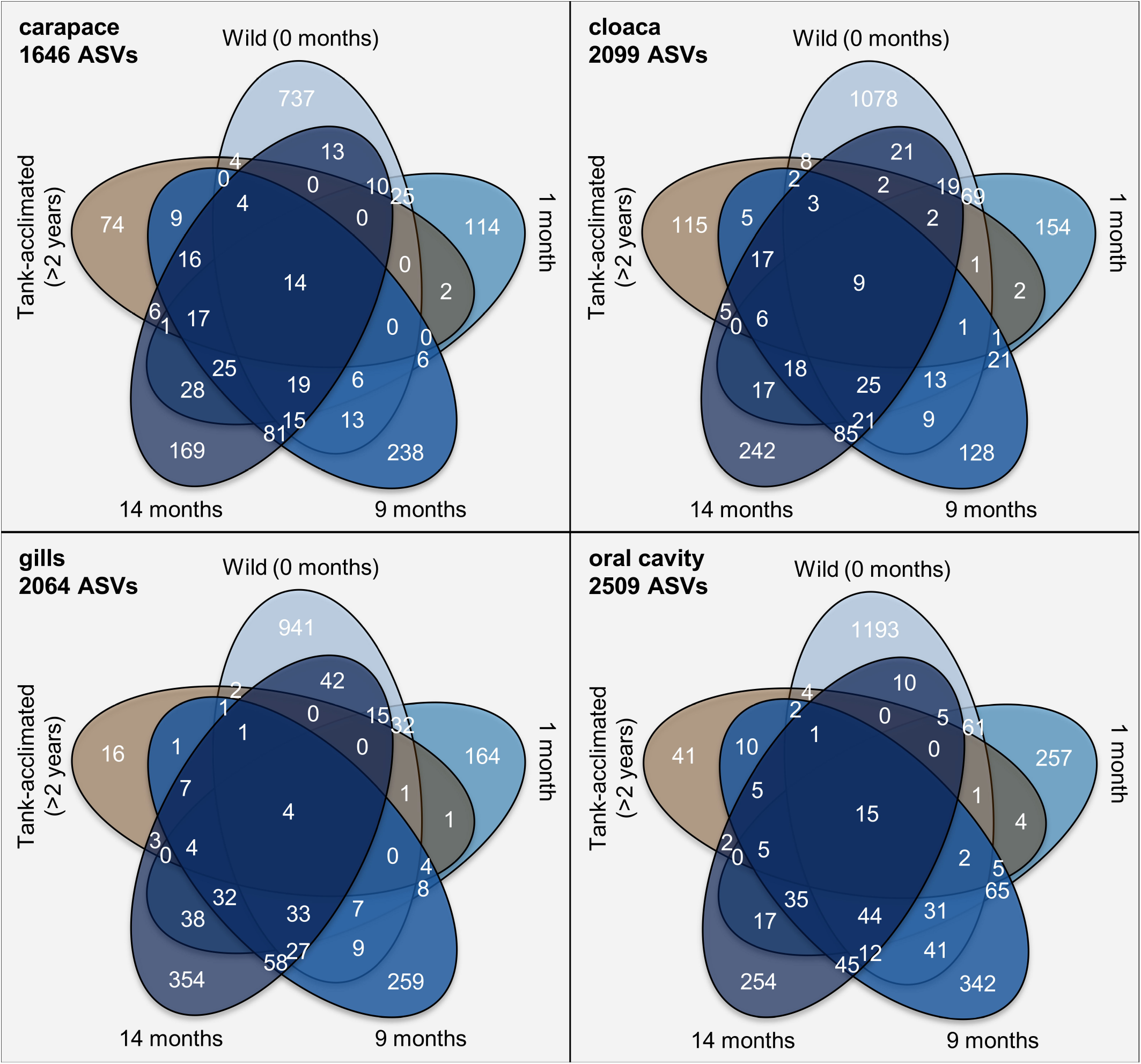
Venn diagrams of the number of amplicon sequence variants (ASVs) identified from different sampling locations on *Limulus polyphemus* over time. In blue, beginning at the top of each diagram is wild-caught prior to captivity (0 months) and moving clock-wise around the diagram are samples from the same individuals at 1 month, 9 months, and 14 months of captivity. The final ellipse is from tank-acclimated individuals (>2 years) sampled at the beginning of the study to establish the microbiome already present in the tank.

Differential abundance analysis was employed to identify ASVs that were significantly (p-value < 0.01) differentially abundant in the wild-caught population between sampling locations or through time in captivity (Figure 5). Several ASVs completely disappeared from the wild horseshoe crab microbiome in captivity, including *Thiothrix*, *Psychrilyobacter*, *Granulosicoccus*, *Alkanindiges,* and uncultivated *Saprospiraceae*. Other ASVs were present in all body locations at the first time point but only partially retained at a lower overall abundance and/or in less body locations after 14 months in captivity, such as *Propionigenium*, *Spiroplasma*, *Pseudoalteromonas*, *Ralstonia*, and *Tenacibaculum*. Additionally, several ASVs associated with *Photobacterium*, *Oleiphilus*, *Spongiimonas*, *Cocleimonas*, *Filomicrobium*, *Arcobacter*, and *Arenicella* either increased in abundance or appeared *de novo* during captivity. Various ASVs associated with unclassified members of *Rhodobacteraceae* were abundant in the wild horseshoe crab microbiome but decreased in abundance and/or prevalence or disappeared completely after time in captivity. After one month in captivity, new ASVs associated with unclassified members of *Rhodobacteraceae* appeared in the wild horseshoe crab microbiome. An additional version of this figure with a more lenient p-value cut-off (p-value = 0.05) is available in the supplementary material (Supplementary Figure 8).

**Figure 5.**
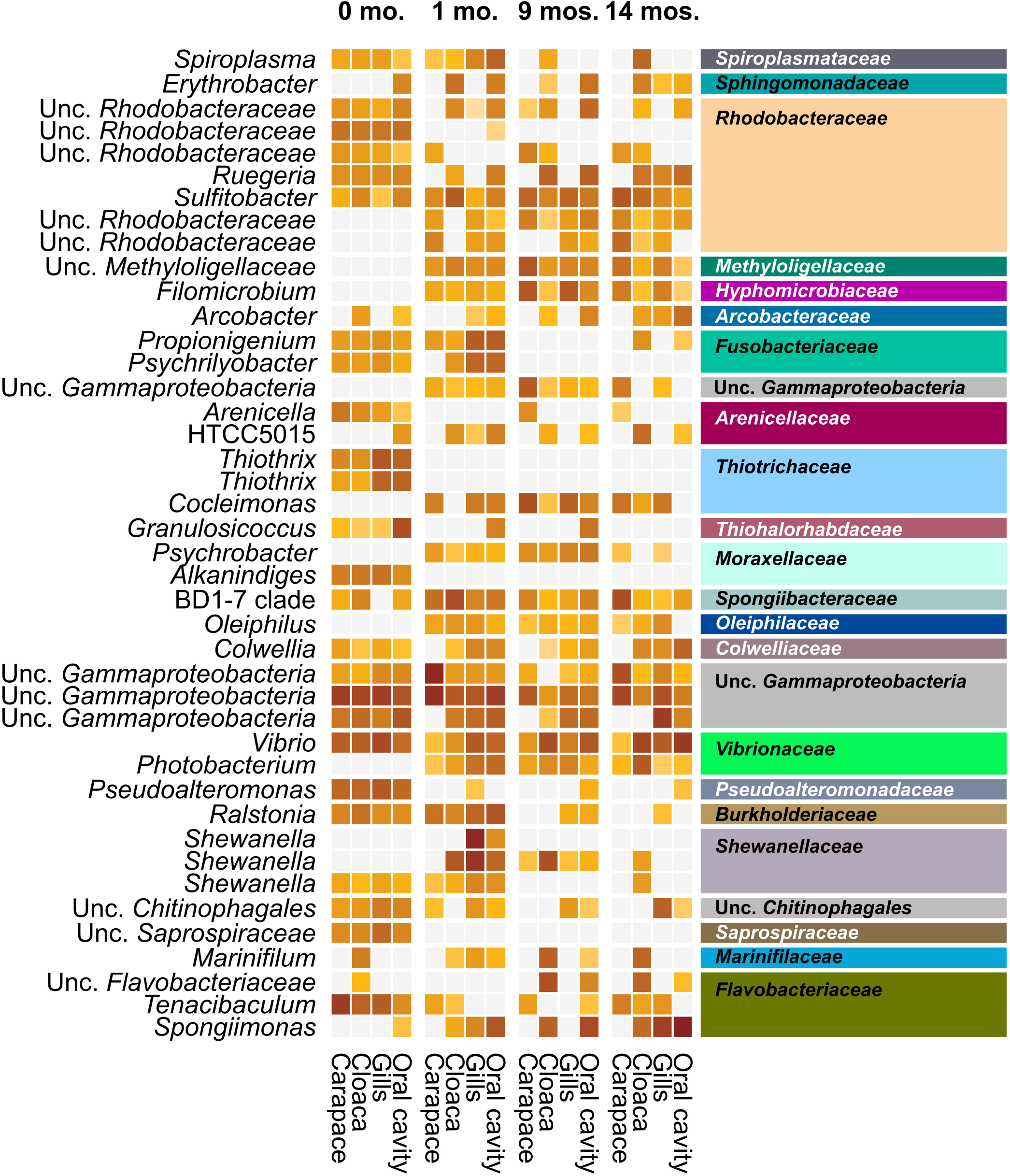
Heatmap showing the log10 counts per million of 16S rRNA gene ASVs deemed significantly differentially abundant across time in captivity in the wild horseshoe crabs via DESeq2 analysis (p-value < 0.01). Samples have been grouped initially by time (0, 1, 9, and 14 months), then by body location (carapace, cloaca, gills, and oral cavity). The tip order of the p < 0.01 ASV tree was used to order the ASVs phylogenetically. ASVs are labeled according to genus-level taxonomy, indicated on the left. Family-level taxonomy of each ASV is indicated on the right side; colors match family-level bar plot (Supplementary Figure 6).

## Discussion

### Environment Plays a Role in Shaping Microbiome of Horseshoe Crabs

The genera *Pseudoalteromonas*, *Vibrio*, and *Photobacterium* have been previously isolated from book gills and mouth samples collected from the wild and lab reared Asian horseshoe crab species *T. gigas* and *C. rotundicauda* (Ismail et al., 2015; Faizul et al., 2015). Although those studies focused on different genera of horseshoe crabs inhabiting geographically distant environments, *Vibrio* and *Pseudoalteromonas* were also detected at appreciable abundance in wild *L. polyphemus*, suggesting they may be part of a natural core horseshoe crab microbiome. *Photobacterium* was also present, however, it was not abundant (<1%) in the microbiomes of wild *L. polyphemus* that we sampled. These three genera are common marine bacteria globally and include known symbionts and commensals of a variety of marine fauna. *Pseudoalteromonas* species are commonly found in the microbiome of different aquatic organisms, including coral species (Shnit-Orland et al., 2012), crayfish (Skelton et al, 2017), and copepods (Moisander et al., 2015). *Vibrio* is commonly found in the microbiomes of aquatic organisms and some species have been identified as pathogens, commensals, and mutualists, yet, an understanding of the relationships between many *Vibrio* species and their hosts requires further research (Thompson et al., 2016). A histological analysis of *Limulus* identified several bacterial species associated with pathological conditions including *Pseudomonas* and *Aeromonas* (LaDouceur et al., 2019) which also appear to be major constituents of what we suspect is a pathological condition in our study, as well. A more comprehensive view of a core horseshoe crab microbiome awaits cultivation-independent surveys of other horseshoe crab species, and more individuals from different geographic and physicochemical environments (e.g., estuarine and marine).

Although little research has been done regarding the microbial communities in horseshoe crabs, extensive work has been carried out for other arthropods. A recent *Proteobacteria*-focused meta-analysis of previous arthropod microbiome studies listed ~500 genera present in the microbiomes of three or more arthropods (Esposti and Romero 2017). Strikingly, only four of these genera were present at high abundance (>5%) in any of the wild horseshoe crabs, possibly reflecting the predominantly terrestrial nature of many arthropods that have been studied and/or the distant phylogenetic relationship between horseshoe crabs and arthropods included in the meta-analysis. These four genera have been described variably in ticks (a fellow arachnid), shrimps, and/or prawns (fellow aquatic arthropods). For example, *Pseudoalteromonas* was prevalent in the microbiomes of wild horseshoe crabs, ticks, and shrimps. *Thiothrix* was seen in horseshoe crabs and ticks but not in the other groups. *Vibrio* was present in all groups but was most abundant in shrimp and ticks. A high abundance of unclassified members of *Rhodobacteraceae* was observed in both horseshoe crabs and shrimp but not in the other two groups, indicating the prevalence of unclassified members of this lineage in marine arthropods. Strikingly, none of the major genera overlapped between scorpions and horseshoe crabs, despite their phylogenetic relationship within the *Arachnida*. Non-proteobacterial members of the wild-caught horseshoe crab microbiome are similarly not commonly reported in terrestrial arthropods (Bouchon et al., 2016; Esposti and Romero 2017; Otani et al., 2014; Vanthournout and Hendrickx 2015).

The meta-analysis validated insights from previous studies that the proteobacterial microbiome of terrestrial arthropods is dominated by taxa commonly found in soil communities, which implies that members of the microbiome are selectively filtered from an environmental pool (Esposti and Romero 2017). By extension, horseshoe crabs could be expected to host predominantly marine and estuarine microorganisms rather than soil microorganisms, but only a few taxa were detected in both the wild-caught horseshoe crab and native Florida substrate samples. However, this is not surprising given the environmental heterogeneity of horseshoe crabs, which provides them with a large pool of environmental microbes to interact with in comparison to our one native substrate sample. Generally, this environmental filtering hypothesis was supported by our data, as we observed many microbes commonly associated with marine systems in the wild horseshoe crab microbiomes. Marine systems are typically dominated by *Alphaproteobacteria*, *Gammaproteobacteria*, and *Bacteroidia* (Louca et al., 2016) and those same three classes often predominate in the microbiomes of marine animals (Colston and Jackson 2016; Esposti and Romero 2017; Thomas et al., 2016).

Some of the microbes in the wild horseshoe crabs reflect their benthic lifestyle and suggest cycling between oxic and anoxic sediments. In particular, the presence of sulfate-reducing bacteria (SRB) (*Desulfobacteraceae* and *Desulfobulbaceae*) on the wild horseshoe crabs is likely enabled by burrowing or foraging in anoxic sediments where SRB predominate; the digestive system of arachnids is permeable to oxygen and not likely to shield SRB from an oxic environment (Lozano-Fernandez et al., 2016). Similarly, the high abundance and diversity of sulfide-oxidizing bacteria (SOB) in the genus *Thiothrix* in wild animals is consistent with the presence of SRB-derived sulfide. Symbioses between sulfur-oxidizing microorganisms and aquatic invertebrates are widespread in sulfide-rich marine environments and thought to have evolved independently in many organisms (Bergin et al., 2017; Dattagupta et al., 2009; Distel et al., 2017; Dubilier et al., 2008). *Thiothrix* is a common ectosymbiont of marine invertebrates (Temara et al., 1992; Brigmon and de Ridder 1998; Bauermeister et al., 2012). The genus *Granulosicoccus* was prevalent in the oral cavity samples of the wild horseshoe crabs and is present in the microbiomes of several other marine organisms, such as seagrass (Kurilenko et al., 2010), kelp (Bengtsson et al., 2012), and corals (van Bleijswijk et al., 2015). A genomic analysis of the type strain *Granulosicoccus antarcticus* IMCC3135T (= KCCM 42676T = NBRC 102684T) revealed the presence of several genes associated with sulfur cycling; for example, the genome contained a gene encoding for the enzyme dimethylsulfoniopropionate (DMSP) demethylase and several genes associated with sulfur oxidation (Kang et al., 2018). These bacteria are likely responsible for a complex sulfur biogeochemical cycle in and on wild horseshoe crabs.

### Significant Shifts Associated with Captivity Observed in Microbial Community Diversity and Structure of Horseshoe Crabs

Public aquaria are popular attractions that are commonplace all around the world as stand-alone entities or as additions to museums, malls, and hotels. Taking advantage of their popularity, modern-day accredited aquaria offer an exciting opportunity for the public to view and interact with various aquatic organisms, while also educating attendees about serious environmental issues, such as conservation. Touch-tank exhibits, which are commonly found in aquaria, provide visitors with the chance to directly interact with and touch live aquatic creatures, such as horseshoe crabs, rays, sharks, and others. Given that aquarium staff members are always present at these exhibits; touch tanks provide a unique educational experience beyond that of just attending an aquarium. Although aquaria and touch-tank exhibits are popular attractions worldwide and serve an important role in furthering ecological education, few studies have explored the health of captive organisms (Johnson et al., 2017; Morris et al., 2012; Persky et al., 2012) or described the microbial communities of the aquarium environment and organisms (Patin et al., 2018), how these microbial communities change over time (Van Bonn et al., 2015; Cardona et al., 2018), and how they compare to the natural environment and wild microbiome (Pagán-Jiménez et al., 2019).

An ever-increasing body of literature has demonstrated the importance of diverse microorganisms in ecology, such as biogeochemical cycling (Rousk and Bengtson 2014), ecosystem health (Hooper et al., 2005), and disease prevention in hosts (Greenspan et al. 2019; Hamdi et al. 2011). In our study, we documented a drastic decrease in the diversity of the wild-caught horseshoe crabs microbiome and significant shifts in microbial community structure shortly after entering captivity. The diversity and composition of the natural microbiome that was lost following entrance into captivity never recovered. The tank-acclimated population, which had spent more than two years in captivity, was extremely dysbiotic. Similar patterns of declines in microbial diversity and microbial community shifts associated with captivity have been observed in sea cucumbers (Pagán-Jiménez et al., 2019) and dugongs (Eigeland et al., 2012). Another study, focused on smooth dogfish, found that there was a high incidence of mortality in adults and pups of an aquarium collection following entrance into captivity, despite various treatments (Persky et al., 2012). Contrastingly, a study assessing the health of cownose rays in a touch-tank exhibit at the John G. Shedd Aquarium to that of a population in an off-exhibit habitat demonstrated that there was no discernible difference in the health of the two populations (Johnson et al., 2017).

Aquarium systems attempt to recreate the natural habitat, but they differ from the natural ecosystem in several ways, including: 1) unnatural physical or chemical conditions (for example, the use of non-native substrate); 2) interactions with organisms that are rarely, if ever, experienced in nature, including microbes seeded from co-habiting species and human contact; 3) an unstable aquarium water microbial community, typically dominated by continuously shifting microbial blooms (Patin et al., 2018; Van Bonn et al., 2015); and 4) increased biomass/population density. We hypothesize that the combined stresses associated with living in a touch-tank exhibit are likely related to the decline in health and microbial diversity of the horseshoe crabs over time in captivity. Similar loss of microbial diversity associated with stress, such as hypoxia or temperature stress, have been described previously in brook char (Boutin et al., 2013) and Pacific oysters (Lokmer and Wegner 2014). Another factor that may have contributed to the microbiome shifts observed during captivity in our study is the difference in diet between wild and captive horseshoe crabs. Wild horseshoe crabs encounter a myriad of possible food sources, including dead fish, algae, mollusks, worms, bivalves, and crustaceans. Captive populations, while still receiving a relatively diverse diet, are not exposed to the same level of diversity as their natural counterparts in the ocean. A captive diet limits not only the variety of food sources available, but also the microbial diversity in food inocula, which has been shown to be important in supporting the development of diverse microbial communities (Bolnick et al., 2014; Heiman and Greenway 2016; Martínez-Mota et al., 2019).

We found that the high-level taxonomy (i.e. phylum and class) of the natural horseshoe crab microbiome was largely retained for over a year, but abundances of individual taxa, such as unclassified members of *Gammaproteobacteria*, *Rhodobacteraceae*, *Flavobacteriaceae*, and *Saprospiraceae,* became increasingly different in all body locations over the course of the experiment. *Thiothrix*, an abundant sulfur-oxidizing bacterium in the wild horseshoe crab microbiome, completely disappeared after only one month in captivity and was replaced by a closely related sulfur-oxidizing genus, *Cocleimonas* (Tanaka et al., 2011). We also observed the appearance of several novel taxa during time in captivity, including, *Spongiimonas*, *Oleiphilus*, *Oleispira*, and unclassified members of *Oceanospirillales* which we hypothesize to be indicative of the captive environment and seeded from the aquarium water (Van Bonn et al., 2015; Patin et al., 2018). Additionally, we recorded several microbial bloom infection events by different taxa during captivity, such as *Shewanella*, *Pseudomonas*, *Cocleimonas*, *Corynebacterium*, *Staphylococcus*, and *Prevotella* 9, suggesting a highly unstable and uneven community following entrance into captivity. Dysbiosis was extreme in the tank-acclimated horseshoe crab microbiomes, which were highly uneven and dominated by *Aeromonas*, *Pseudomonas*, and unclassified *Enterobacteriaceae*, with *Shewanella* and *Enterococcus* also abundant in the cloaca. The appearance of human-associated taxa, such as *Enterococcus*, *Corynebacterium*, *Staphylococcus*, and *Prevotella*, in the captive animal populations may be related to the dynamic nature of the touch-tank exhibit, where various visitors are in direct contact with the animals. This might facilitate the transfer of human-associated taxa to the captive horseshoe crabs. This is in direct opposition to a study on cownose rays in touch tanks, which concluded that the transfer of human-associated taxa to the rays was negligible (Kearns et al., 2017).

### Symbiotic Potential of Taxa Identified in the Wild Horseshoe Crab Microbiome

Termites and cockroaches are well-studied arthropod model organisms in symbiosis research and several studies have demonstrated the importance that symbiotic interactions played in their evolution, particularly their expansion into previously unoccupied niches, such as plant polymer degradation (Berlanga and Guerrero 2016). In addition, marine symbioses are also quite well researched, with various studies detailing the biotechnological potential of and interactions between host-symbiont in deep-sea hydrothermal vents (Ho et al. 2017; Zimmermann et al. 2014), coral reefs (Venn et al., 2008), sponges (Venn et al., 2008), and polychaetes (Goffredi et al. 2005). We speculate that sulfur-cycling microorganisms, such as *Thiothrix* and *Granulosicoccus*, may play an important role in mediating horseshoe crab health and microbiome composition, possibly by competing with pathogens for attachment space and/or nutrients. As discussed previously, these microorganisms would benefit by living at or near redox interfaces and/or by transitioning frequently between oxic and anoxic environments that may source terminal electron acceptors and donors, respectively, and a variety of other nutrients.

One pathology associated with captive horseshoe crabs is the appearance of pitting or lesions on the carapace, which is composed primarily of chitin. Once the exoskeleton of the horseshoe crab has been breached (or degraded), it would then be susceptible to secondary infections and given that adults do not molt, the damage accumulated is irreparable. Previous studies have associated infections in captive lab horseshoe crab populations with eukaryotic parasites, algae, fungi, and bacteria (commonly *Cyanobacteria* and Gram-negative bacteria) (Leibovitz and Lewbart 2004, Smith 2006; Braverman et al., 2012, LaDouceur et al, 2019). We hypothesize that the appearance of these lesions could be due to dysbiosis of the horseshoe crab microbiome, resulting in an overall increase in the abundance of chitinolytic bacteria or the appearance of opportunistic pathogens capable of degrading chitin. More specifically, we posit that certain taxa in the wild animals with potential chitinolytic activity, such as *Vibrio* and *Pseudoalteromonas* (Machado et al., 2015), may represent commensalistic or mutualistic members of the microbiome that help remove loose material, while out-competing potentially pathogenic chitinolytic bacteria. These organisms are also likely kept in check by other microorganisms in nature. In contrast, we observed blooms of potentially pathogenic *Aeromonas* and *Pseudomonas* ASVs in captivity, which are likely to produce lesions in the context of such a dysbiosis (Frederiksen et al. 2013). Additionally, several species of *Pseudoalteromonas* are widely discussed in the literature as symbionts, given their diverse metabolic potential and their ability to produce a variety of antibacterial, antifungal, algicidal, and antifouling compounds (Atencio et al. 2018; Offret et al. 2016). Fungi have been identified as possible chitinase containing pathogens that could be involved in this process (Tuxbury et al., 2014; LaDouceur et al., 2019). In our study, we excluded eukaryotic sequences from analysis and therefore were unable to document the fungal communities of our samples.

## Concluding Remarks

This study details the first cultivation-independent survey of the horseshoe crab microbiome. The microbial community of wild horseshoe crabs was diverse and highly similar across body locations but was nearly completely restructured through time in captivity. Changes included significant loss of diversity, increasing uniqueness by body location, transient microbial bloom events, and eventually development of highly uneven dysbiotic communities dominated by a few opportunistic pathogens, primarily *Aeromonas* and *Pseudomonas*.

This study provides an initial framework for understanding the horseshoe crab microbiome and its response to captivity. We suggest some directions for future study below. Sampling could be increased and expanded. The wild horseshoe crabs that were sampled represent a very small portion of the geographic distribution and physicochemical conditions of *L. polyphemus* in the wild. To more deeply understand the diversity of the natural microbiome of these animals, additional sampling throughout their range and at different points in their life cycle is warranted. A longer experimental timeline (> two years in captivity) and more frequent sampling could provide a more detailed view of how the microbiota change in captivity. While we found evidence of bacteria that are known to contain chitinase, it is still unknown if the carapace lesions are caused by bacteria or if the lesions develop first, then become opportunistically infected after. A directed investigation that includes progressively analyzing the lesions as they develop or intentionally infecting individuals with different strains of bacteria could shed light on the mechanism of disease. Several bacteria found on captive horseshoe crabs were typical of human hosts and others were known human pathogens—understanding more about the transmission of these bacteria in both directions is important for the health and safety of both the animals and the visitors of the exhibit. Other avenues to explore include understanding the role substrate type and depth plays in microbial associations and how the density of individuals including co-habiting species affect the rate at which the microbial community changes.

## Author Contributions

SAN, JJ, and LRB conceived of the study and experimental design. Samples were collected and processed by SAN, LRB, FL, GM, AF, and CS. SN, BPH, AF, and CS worked together to design the figures, but AF and CS were responsible for generating figures for the manuscript. AF and SAN co-wrote the first draft of this manuscript, which was initially edited by BPH. Subsequent drafts were edited by all authors.

## Acknowledgments

We would like to thank Nicole Cox, and Bianca Markie for their excellent care of the animals and assistance with sampling during this study.

## Supplementary figure legends

**Figure 1.** Rarefaction curve comparing the number of total observed species (Species) to the number of reads in each sample (Sample Size). Samples are colored corresponding to the time series: substrates (pink), wild (shades of blue), and tank-acclimated (brown).

**Figure 2.** Bar plot representing the domain-level relative abundance of taxa in all samples. The two substrate samples have been grouped and all horseshoe crabs have been grouped by individual sampled. Taxonomic classification was based on SILVA (version 132).

**Figure 3.** Bar plot representing the phylum-level relative abundance of taxa in all samples. The two substrate samples have been grouped and all horseshoe crabs have been grouped by individual sampled. Taxa whose abundance was <5% were grouped together and listed as such. All unclassified taxa at this rank and the <5% group are colored gray. Taxonomic classification was based on SILVA (version 132).

**Figure 4.** Bar plot representing the class-level relative abundance of taxa in all samples. The two substrate samples have been grouped and all horseshoe crabs have been grouped by individual sampled. Taxa whose abundance was <5% were grouped together and listed as such. All unclassified taxa at this rank and the <5% group are colored gray. Taxonomic classification was based on SILVA (version 132).

**Figure 5.** Bar plot representing the order-level relative abundance of taxa in all samples. The two substrate samples have been grouped and all horseshoe crabs have been grouped by individual sampled. Taxa whose abundance was <5% were grouped together and listed as such. All unclassified taxa at this rank and the <5% group are colored gray. Taxonomic classification was based on SILVA (version 132).

**Figure 6.** Bar plot representing the family-level relative abundance of taxa in all samples. The two substrate samples have been grouped and all horseshoe crabs have been grouped by individual sampled. Taxa whose abundance was <5% were grouped together and listed as such. All unclassified taxa at this rank and the <5% group are colored gray. Taxonomic classification was based on SILVA (version 132).

**Figure 7.** Bar plot representing the genus-level relative abundance of taxa in all samples. The two substrate samples have been grouped and all horseshoe crabs have been grouped by individual sampled. Taxa whose abundance was <5% were grouped together and listed as such. All unclassified taxa at this rank and the <5% group are colored gray. Taxonomic classification was based on SILVA (version 132).

**Figure 8.** Heatmap showing the log10 counts per million of 16S rRNA gene ASVs deemed significantly differentially abundant across time in captivity in the wild horseshoe crabs via DESeq2 analysis (p < 0.05). Samples have been grouped initially by time (0, 1, 9, and 14 months), then by body location (carapace, cloaca, gills, and oral cavity). The tip order of the p < 0.05 ASV tree was used to order the ASVs phylogenetically. ASVs are labeled according to Genus-level taxonomy, indicated on the left. Family-level taxonomy of each ASV is indicated on the right side; colors match family-level bar plot (Supplementary Figure 6).

## References

Atencio, L. A., Dal Grande, F., Young, G. O., Gavilán, R., Guzmán, H. M., Schmitt, I., Mejía, L., and Gutiérrez, M. (2018). Antimicrobial-producing *Pseudoalteromonas* from the marine environment of Panama shows a high phylogenetic diversity and clonal structure. J. Basic Microbiol. 58, 747–769. doi: 10.1002/jobm.201800087

Ballesteros, J.A. and Sharma, P.P. (2019). A critical appraisal of the placement of *Xiphosura* (*Chelicerata*) with account of known sources of phylogenetic error. Syst. Biol. 68, 896–917. doi: 10.1093/sysbio/syz011

Bauermeister, J., Ramette, A. and Dattagupta, S. (2012). Repeatedly evolved host-specific ectosymbioses between sulfur-oxidizing bacteria and amphipods living in a cave ecosystem. PLoS ONE 7:e50254. doi: 10.1371/journal.pone.0050254

Bengtsson, M. M., Sjøtun, K., Lanzén, A., & Øvreås, L. (2012). Bacterial diversity in relation to secondary production and succession on surfaces of the kelp *Laminaria hyperborea*. ISME J. 6, 2188–2198. doi: 10.1038/ismej.2012.67

Bergin, C., Wentrup, C., Brewig, N., Blazejak, A., Erséus, C., Giere, O., Schmid, M., De Wit, P. and Dubilier, N. (2018). Acquisition of a novel sulfur-oxidizing symbiont in the gutless marine worm *Inanidrilus exumae*. Appl. Environ. Microbiol. 84:e02267–17. doi: 10.1128/AEM.02267-17

Berlanga, M. and Guerrero, R. (2016). The holobiont concept: the case of xylophagous termites and cockroaches. Symbiosis 68, 49–60. doi: 10.1007/s13199-016-0388-9

van Bleijswijk, J. D. L., Whalen, C., Duineveld, G. C. A., Lavaleye, M. S. S., Witte, H. J., & Mienis, F. (2015). Microbial assemblages on a cold-water coral mound at the SE Rockall Bank (NE Atlantic): interactions with hydrography and topography. Biogeosciences 12, 4483–4496. doi: 10.5194/bg-12-4483-2015

Bokulich, N.A., Kaehler, B.D., Rideout, J.R., Dillon, M., Bolyen, E., Knight, R., Huttley, G.A. and Caporaso, J.G. (2018). Optimizing taxonomic classification of marker-gene amplicon sequences with QIIME 2’s q2-feature-classifier plugin. Microbiome 6, 90. doi: 10.1186/s40168-018-0470-z

Bolnick, D. I., Snowberg, L. K., Hirsch, P. E., Lauber, C. L., Knight, R., Caporaso, J. G., and Svanbäck, R. (2014). Individuals’ diet diversity influences gut microbial diversity in two freshwater fish (threespine stickleback and Eurasian perch). Ecol. Lett. 17, 979–987. doi: 10.1111/ele.12301

van Bonn, W., LaPointe, A., Gibbons, S.M., Frazier, A., Hampton□Marcell, J. and Gilbert, J. (2015). Aquarium microbiome response to ninety□percent system water change: Clues to microbiome management. Zoo Biol. 34, 360–367. doi: 10.1002/zoo.21220

Bouchon, D., Zimmer, M. and Dittmer, J. (2016). The terrestrial isopod microbiome: an all-in-one toolbox for animal–microbe interactions of ecological relevance. Front. Microbiol. 7:1472. doi: 10.3389/fmicb.2016.01472

Boutin, S., Bernatchez, L., Audet, C. and Derôme, N. (2013). Network analysis highlights complex interactions between pathogen, host and commensal microbiota. PLoS ONE 8:e84772. doi: 10.1371/journal.pone.0084772

Braverman, H., Leibovitz, L. and Lewbart, G.A. (2012). Green algal infection of American horseshoe crab (*Limulus polyphemus*) exoskeletal structures. J. Invertebr. Pathol. 111, 90–93. doi: 10.1016/j.jip.2012.06.002

Brigmon, R.L. and De Ridder, C. (1998). Symbiotic relationship of *Thiothrix* spp. with an echinoderm. Appl. Environ. Microbiol. 64, 3491–3495.

Callahan, B. J., McMurdie, P. J., Rosen, M. J., Han, A. W., Johnson, A. J. A., and Holmes, S. P. (2016). DADA2: high-resolution sample inference from Illumina amplicon data. Nat. Methods 13, 581–583. doi: 10.1038/nmeth.3869

Caporaso, J.G., Kuczynski, J., Stombaugh, J., Bittinger, K., Bushman, F.D., Costello, E.K., Fierer, N., Pena, A.G., Goodrich, J.K., Gordon, J.I. and Huttley, G.A. (2010). QIIME allows analysis of high-throughput community sequencing data. Nat. Methods 7:335–336. doi: 10.1038/nmeth.f.303

Cardona, C., Lax, S., Larsen, P., Stephens, B., Hampton-Marcell, J., Edwardson, C.F., Henry, C., Van Bonn, B. and Gilbert, J.A. (2018). Environmental sources of bacteria differentially influence host-associated microbial dynamics. MSystems 3:e00052–18. doi: 10.1128/mSystems.00052-18

Carmichael, R. H. and Brush, E. (2012). Three decades of horseshoe crab rearing: a review of conditions for captive growth and survival. Rev. Aquacult. 4, 32–43.

Colston T.J. and Jackson C.R. (2016) Microbiome evolution along divergent branches of the vertebrate tree of life: what is known and unknown. Mol Ecol. 25: 3776–3800. doi: 10.1111/mec.13730

Dattagupta, S., Schaperdoth, I., Montanari, A., Mariani, S., Kita, N., Valley, J.W. and Macalady, J.L. (2009). A novel symbiosis between chemoautotrophic bacteria and a freshwater cave amphipod. ISME J. 3, 935–943. doi: 10.1038/ismej.2009.34

Distel, D.L., Altamia, M.A., Lin, Z., Shipway, J.R., Han, A., Forteza, I., Antemano, R., Limbaco, M.G.J.P., Tebo, A.G., Dechavez, R. and Albano, J. (2017). Discovery of chemoautotrophic symbiosis in the giant shipworm *Kuphus polythalamia* (*Bivalvia*: *Teredinidae*) extends wooden-steps theory. Proc. Natl. Acad. Sci. U.S.A. 114, 3652–3658. doi: 10.1073/pnas.1620470114

Dixon, P. (2003). VEGAN, a package of R functions for community ecology. J. Vegetation Sci. 14, 927–930. doi: 10.1111/j.1654-1103.2003.tb02228.x

Dubilier, N., Bergin, C. and Lott, C. (2008). Symbiotic diversity in marine animals: the art of harnessing chemosynthesis. Nat. Rev. Microbiol. 6, 725–740. doi: 10.1038/nrmicro1992

Eigeland, K. A., Lanyon, J. M., Trott, D. J., Ouwerkerk, D., Blanshard, W., Milinovich, G. J., … & Klieve, A. V. (2012). Bacterial community structure in the hindgut of wild and captive dugongs (Dugong dugon). Aquat. Mamm. 38, 402–411. doi: 10.1578/AM.38.4.2012.402

Frederiksen, R.F., Paspaliari, D.K., Larsen, T., Storgaard, B.G., Larsen, M.H., Ingmer, H., Palcic, M.M. and Leisner, J.J. (2013). Bacterial chitinases and chitin-binding proteins as virulence factors. Microbiology 159, 833–847. doi: 10.1099/mic.0.051839-0

Goffredi, S.K., Orphan, V.J., Rouse, G.W., Jahnke, L., Embaye, T., Turk, K., Lee, R. and Vrijenhoek, R.C. (2005). Evolutionary innovation: a bone□eating marine symbiosis. Environ. Microbiol. 7, 1369–1378. doi: 10.1111/j.1462-2920.2005.00824.x

Greenspan, S.E., Lyra, M.L., Migliorini, G.H., Kersch-Becker, M.F., Bletz, M.C., Lisboa, C.S., Pontes, M.R., Ribeiro, L.P., Neely, W.J., Rezende, F. and Romero, G.Q. (2019). Arthropod–bacteria interactions influence assembly of aquatic host microbiome and pathogen defense. P. Roy. Soc. B-Biol. Sci. 286:20190924. doi: 10.1098/rspb.2019.0924

Hamdi, C., Balloi, A., Essanaa, J., Crotti, E., Gonella, E., Raddadi, N., Ricci, I., Boudabous, A., Borin, S., Manino, A. and Bandi, C. (2011). Gut microbiome dysbiosis and honeybee health. J. Appl. Entomol. 135, 524–533. doi: 10.1111/j.1439-0418.2010.01609.x

Hargreaves, J.A. (1998). Nitrogen biogeochemistry of aquaculture ponds. Aquaculture 166, 181–212.

Heiman, M. L. and Greenway, F. L. (2016). A healthy gastrointestinal microbiome is dependent on dietary diversity. Mol. Metab. 5, 317–320. doi: 10.1016/j.molmet.2016.02.005

Ho, P.T., Park, E., Hong, S.G., Kim, E.H., Kim, K., Jang, S.J., Vrijenhoek, R.C. and Won, Y.J. (2017). Geographical structure of endosymbiotic bacteria hosted by *Bathymodiolus* mussels at eastern Pacific hydrothermal vents. BMC Evol. Biol. 17: 121. doi: 10.1186/s12862-017-0966-3

Hooper, D.U., Chapin, F.S., Ewel, J.J., Hector, A., Inchausti, P., Lavorel, S., Lawton, J.H., Lodge, D.M., Loreau, M., Naeem, S. and Schmid, B. (2005). Effects of biodiversity on ecosystem functioning: a consensus of current knowledge. Ecol. Monogr. 75, 3–35. doi: 10.1890/04-0922

Ismail, N., Faridah, M., Ahmad, A., Alia’m, A. A., Khai, O. S., Sofa, M. F. A. M., & Manca, A. (2015). Marine Bacteria Associated with Horseshoe Crabs, *Tachypleus gigas* and *Carcinoscorpius rotundicauda*. In Changing Global Perspectives on Horseshoe Crab Biology, Conservation and Management (pp. 313–320). Springer, Cham. doi: 10.1007/978-3-319-19542-1_18

Johnson, J.G., Naples, L.M., Van Bonn, W.G., Kent, A.D., Mitchell, M.A. and Allender, M.C. (2017). Evaluation of health parameters in cownose rays (*Rhinoptera bonasus*) housed in a seasonal touch pool habitat compared with an off-exhibit habitat. J. Zoo Wildlife Med. 48, 954–960. doi: 10.1638/2017-0091.1

Kalyaanamoorthy, S., Minh, B. Q., Wong, T. K. F., Haeseler, A. von, & Jermiin, L. S. (2017). ModelFinder: Fast model selection for accurate phylogenetic estimates. Nat. Methods 14, 587–589. https://doi.org/10.1038/nmeth.4285

Kang, I., Lim, Y., & Cho, J. C. (2018). Complete genome sequence of *Granulosicoccus antarcticus* type strain IMCC3135T, a marine gammaproteobacterium with a putative dimethylsulfoniopropionate demethylase gene. Mar. Genomics 37, 176–181. doi: 10.1016/j.margen.2017.11.005

Kearns, P. J., Bowen, J. L., & Tlusty, M. F. (2017). The skin microbiome of cow□nose rays (Rhinoptera bonasus) in an aquarium touch□tank exhibit. Zoo Biol. 36, 226–230. doi: 10.1002/zoo.21362

Kurilenko, V. V., Christen, R., Zhukova, N. V., Kalinovskaya, N. I., Mikhailov, V. V., Crawford, R. J., & Ivanova, E. P. (2010). *Granulosicoccus coccoides* sp. nov., isolated from leaves of seagrass (*Zostera marina*). Int. J. Syst. Evol. Microbiol. 60, 972–976. doi: 10.1099/ijs.0.013516-0

Leibovitz, L. (1986). Cyanobacterial diseases of the horseshoe crab (*Limulus polyphemus*). Bio. Bull. 171, 482–483).

LaDouceur, E. E. B., Mangus, L., Garner, M. M., & Cartoceti, A. N. (2019). Histologic findings in captive American horseshoe crabs (*Limulus polyphemus*). Vet. Pathol. 6, 932–939. doi: 10.1177/0300985819859877

Liu, J. S., and Passaglia, C. L. (2009). Using the horseshoe crab, *Limulus polyphemus*, in vision research. J. Vis. Exp. 29:e1384.

Llewellyn, M. S., Boutin, S., Hoseinifar, S. H., & Derome, N. (2014). Teleost microbiomes: the state of the art in their characterization, manipulation and importance in aquaculture and fisheries. Front. Microbiol. 5; 207. doi: 10.3389/fmicb.2014.00207

Lokmer, A. and Wegner, K.M. (2015). Hemolymph microbiome of Pacific oysters in response to temperature, temperature lstress and infection. ISME J. 9:670. doi: 10.1038/ismej.2014.160

Love, M.I., Huber, W. and Anders, S. (2014). Moderated estimation of fold change and dispersion for RNA-seq data with DESeq2. Genome Biol. 15:550. doi: 10.1186/s13059-014-0550-8

Lozano-Fernandez J., Carton R., Tanner A.R., Puttick M.N., Blaxter M., Vinther J., et al. (2016). A molecular palaeobiological exploration of arthropod terrestrialization. Philos. Trans. R. Soc. Lond. B Biol Sci. 371, 20150133. doi: 10.1098/rstb.2015.0133

Machado, H., Sonnenschein, E.C., Melchiorsen, J. and Gram, L. (2015). Genome mining reveals unlocked bioactive potential of marine Gram-negative bacteria. BMC Genomics 16:158. doi: 10.1186/s12864-015-1365-z

Martínez-Mota, R., Kohl, K. D., Orr, T. J., and Dearing, M. D. (2020). Natural diets promote retention of the native gut microbiota in captive rodents. ISME J. 14, 67–78. doi: 10.1038/s41396-019-0497-6

McMurdie, P.J. and Holmes, S. (2013). phyloseq: an R package for reproducible interactive analysis and graphics of microbiome census data. PLoS ONE 8:e61217. doi: 10.1371/journal.pone.0061217

Minh, B. Q., Nguyen, M. A. T., & von Haeseler, A. (2013). Ultrafast Approximation for Phylogenetic Bootstrap. Mol. Biol. Evol. 30, 1188–1195. doi: 10.1093/molbev/mst024

Moisander, P.H., Sexton, A.D. and Daley, M.C. (2015). Stable associations masked by temporal variability in the marine copepod microbiome. PLoS ONE 10:e0138967. doi: 10.1371/journal.pone.0138967

Morris, A.L., Stremme, D.W., Sheppard, B.J., Walsh, M.T., Farina, L.L. and Francis-Floyd, R. (2012). The onset of goiter in several species of sharks following the addition of ozone to a touch pool. J. Zoo Wildlife Med. 621–624.

Nguyen, L.T., Schmidt, H.A., von Haeseler, A., & Minh, B.Q. (2015). IQ-TREE: A Fast and Effective Stochastic Algorithm for Estimating Maximum-Likelihood Phylogenies. Mol. Biol. Evol. 32, 268–274. doi: 10.1093/molbev/msu300

Nolan, M.W. and Smith, S.A. (2009). Clinical evaluation, common diseases, and veterinary care of the horseshoe crab, *Limulus polyphemus*. In Biology and Conservation of Horseshoe Crabs, 479–499. Springer, Boston, MA. doi: 10.1007/978-0-387-89959-6_30

Offret, C., Desriac, F., Le Chevalier, P., Mounier, J., Jégou, C., and Fleury, Y. (2016). Spotlight on antimicrobial metabolites from the marine bacteria *Pseudoalteromonas*: Chemodiversity and ecological significance. Mar. Drugs 14:129. doi: 10.3390/md14070129

Otani, S., Mikaelyan, A., Nobre, T., Hansen, L.H., Koné, N.G.A., Sørensen, S.J., Aanen, D.K., Boomsma, J.J., Brune, A. and Poulsen, M. (2014). Identifying the core microbial community in the gut of fungus□growing termites. Mol. Ecol. 23, 4631–4644. doi: 10.1111/mec.12874

Pagán-Jiménez, M., Ruiz-Calderón, J.F., Dominguez-Bello, M.G. and García-Arrarás, J.E. (2019). Characterization of the intestinal microbiota of the sea cucumber *Holothuria glaberrima*. PLoS ONE 14:e0208011. doi: 10.1371/journal.pone.0208011

Patin, N.V., Pratte, Z.A., Regensburger, M., Hall, E., Gilde, K., Dove, A.D. and Stewart, F.J. (2018). Microbiome dynamics in a large artificial seawater aquarium. Appl. Environ. Microbiol. 84, 00179–18. doi: 10.1128/AEM.00179-18

Paradis, E., Claude, J., & Strimmer, K. (2004). APE: Analyses of Phylogenetics and Evolution in R language. Bioinformatics 20, 289–290. doi: 10.1093/bioinformatics/btg412

Persky, M.E., Williams, J.J., Burks, R.E., Bowman, M.R., Ramer, J.C. and Proudfoot, J.S. (2012). Hematologic, plasma biochemistry, and select nutrient values in captive smooth dogfish (*Mustelus canis*). J. Zoo Wildlife Med., 842–851.

Price, M.N., Dehal, P.S. and Arkin, A.P. (2009). FastTree: computing large minimum evolution trees with profiles instead of a distance matrix. Mol. Biol. Evol. 26, 1641–1650. doi: 10.1093/molbev/msp077

Price, M.N., Dehal, P.S. and Arkin, A.P. (2010). FastTree 2–approximately maximum-likelihood trees for large alignments. PLoS ONE 5:e9490. doi: 10.1371/journal.pone.0009490

Pruesse, E., Quast, C., Knittel, K., Fuchs, B.M., Ludwig, W., Peplies, J. and Glöckner, F.O. (2007). SILVA: a comprehensive online resource for quality checked and aligned ribosomal RNA sequence data compatible with ARB. Nucleic Acids Res. 35, 7188–7196. doi: 10.1093/nar/gkm864

Pruesse, E., Peplies, J., & Glöckner, F. O. (2012). SINA: Accurate high-throughput multiple sequence alignment of ribosomal RNA genes. Bioinformatics 28, 1823–1829. doi: 10.1093/bioinformatics/bts252

Revell, L.J. (2012). phytools: An R package for phylogenetic comparative biology (and other things). Methods in Ecol. and Evol. 3, 217–223. https://doi.org/10.1111/j.2041-210X.2011.00169.x

Rousk, J. and Bengtson, P. (2014). Microbial regulation of global biogeochemical cycles. Front. Microbiol. 5:103. doi: 10.3389/fmicb.2014.00103

Rudkin, D. M., & Young, G. A. (2009). Horseshoe crabs–an ancient ancestry revealed. In Biology and conservation of horseshoe crabs, 25-44. Springer, Boston, MA. doi: 10.1007/978-0-387-89959-6_2

Shinn, A.P., Mühlhölzl, A.P., Coates, C.J., Metochis, C., and Freeman, M.A. (2015) *Zoothamnium duplicatum* infestation of cultured horseshoe crabs (*Limulus polyphemus*). J. Invertebr. Pathol. 125, 81–86. doi: 10.1016/j.jip.2014.12.002

Shnit-Orland, M., Sivan, A. and Kushmaro, A. (2012). Antibacterial activity of *Pseudoalteromonas* in the coral holobiont. Microb. Ecol. 64, 851–859. doi: 10.1007/s00248-012-0086-y

Skelton, J., Geyer, K.M., Lennon, J.T., Creed, R.P. and Brown, B.L. (2017). Multi-scale ecological filters shape the crayfish microbiome. Symbiosis 72, 159–170. doi: 10.1007/s13199-016-0469-9

Smith, D.R. (2007). Effect of horseshoe crab spawning density on nest disturbance and exhumation of eggs: a simulation study. Estuar. Coast. 30, 287–295. doi: 10.1007/BF02700171

Smith, D.R., Brockmann, H.J., Beekey, M.A., King, T.L., Millard, M.J., & Zaldívar-rae, J. (2017). Conservation status of the American horseshoe crab, (*Limulus polyphemus*): A regional assessment. Rev. Fish Biol. and Fisher. 1, 135–175. doi: 10.1007/s11160-016-9461-y

Tanaka, N., Romanenko, L. A., Iino, T., Frolova, G. M., & Mikhailov, V. V. (2011). *Cocleimonas flava* gen. nov., sp. nov., a gammaproteobacterium isolated from sand snail (*Umbonium costatum*). Int. J. Syst. Evol. Microbiol. 61, 412–416. doi: 10.1099/ijs.0.020263-0

Temara, A., De Ridder, C., Kuenen, J.G. and Robertson, L.A. (1993). Sulfide-oxidizing bacteria in the burrowing echinoid, *Echinocardium cordatum* (*Echinodermata*). Mar. Biol. 115, 179–185. doi: 10.1007/BF00346333

Thomas T., Moitinho-Silva L., Lurgi M., Björk J.R., Easson C., Astudillo-García C., et al. (2016). Diversity, structure and convergent evolution of the global sponge microbiome. Nat. Commun. 7: 11870. doi:10.1038/ncomms11870

Thompson, J., Ben Ahmeid, A.A., Pickett, C. and Neil, D. (2011). Identification of cost-effective measures to improve holding conditions and husbandry practices for the horseshoe crab *Limulus polyphemus*. Project Report. University of Glasgow, Glasgow, UK

Thompson, F., Iida, T., and Swings, J. (2016). Biodiversity of Vibrios. Microbiol. Mol. Biol. R. 68, 403–431. doi: 10.1128/MMBR.68.3.403-431.2004

Thompson, L.R., Sanders, J.G., McDonald, D., Amir, A., Ladau, J., Locey, K.J., Prill, R.J., Tripathi, A., Gibbons, S.M., Ackermann, G. and Navas-Molina, J.A. (2017) A communal catalogue reveals Earth’s multiscale microbial diversity. Nature 55, 457–463. doi: 10.1038/nature24621

Tuxbury, K.A., Shaw, G.C., Montali, R.J., Clayton, L.A., Kwiatkowski, N.P., Dykstra, M.J., Mankowski, J.L. (2014) *Fusarium solani* species complex associated with carapace lesions and branchitis in captive American horseshoe crabs *Limulus polyphemus*. Dis Aquat Org 109, 223–230. doi: 10.3354/dao02764

Vanthournout, B. and Hendrickx, F. (2015). Endosymbiont dominated bacterial communities in a dwarf spider. PLoS ONE 10:0117297. doi: 10.1371/journal.pone.0117297

Venn, A.A., Loram, J.E. and Douglas, A.E. (2008). Photosynthetic symbioses in animals. J. Exp. Bot. 59, 1069–1080. doi: 10.1093/jxb/erm328

Wickham, H. (2011). Ggplot2. WIREs Comp. Stat. 2, 180–185. doi: 10.1002/wics.147

Williams K.L. (2019) *Limulus* as a Model Organism. In Endotoxin Detection and Control in Pharma, Limulus, and Mammalian Systems, 597–629. Springer, Cham

Zimmermann, J., Lott, C., Weber, M., Ramette, A., Bright, M., Dubilier, N. and Petersen, J.M. (2014). Dual symbiosis with co□occurring sulfur□oxidizing symbionts in vestimentiferan tubeworms from a Mediterranean hydrothermal vent. Environ. Microbiol. 16, 3638–3656. doi: 10.1111/1462-2920.12427

