## Supplementary figures and images for "Microbiome shifts associated with the introduction of wild Atlantic horseshoe crabs (*Limulus polyphemus*) into a touch-tank exhibit"

### Supplemental Figure 1

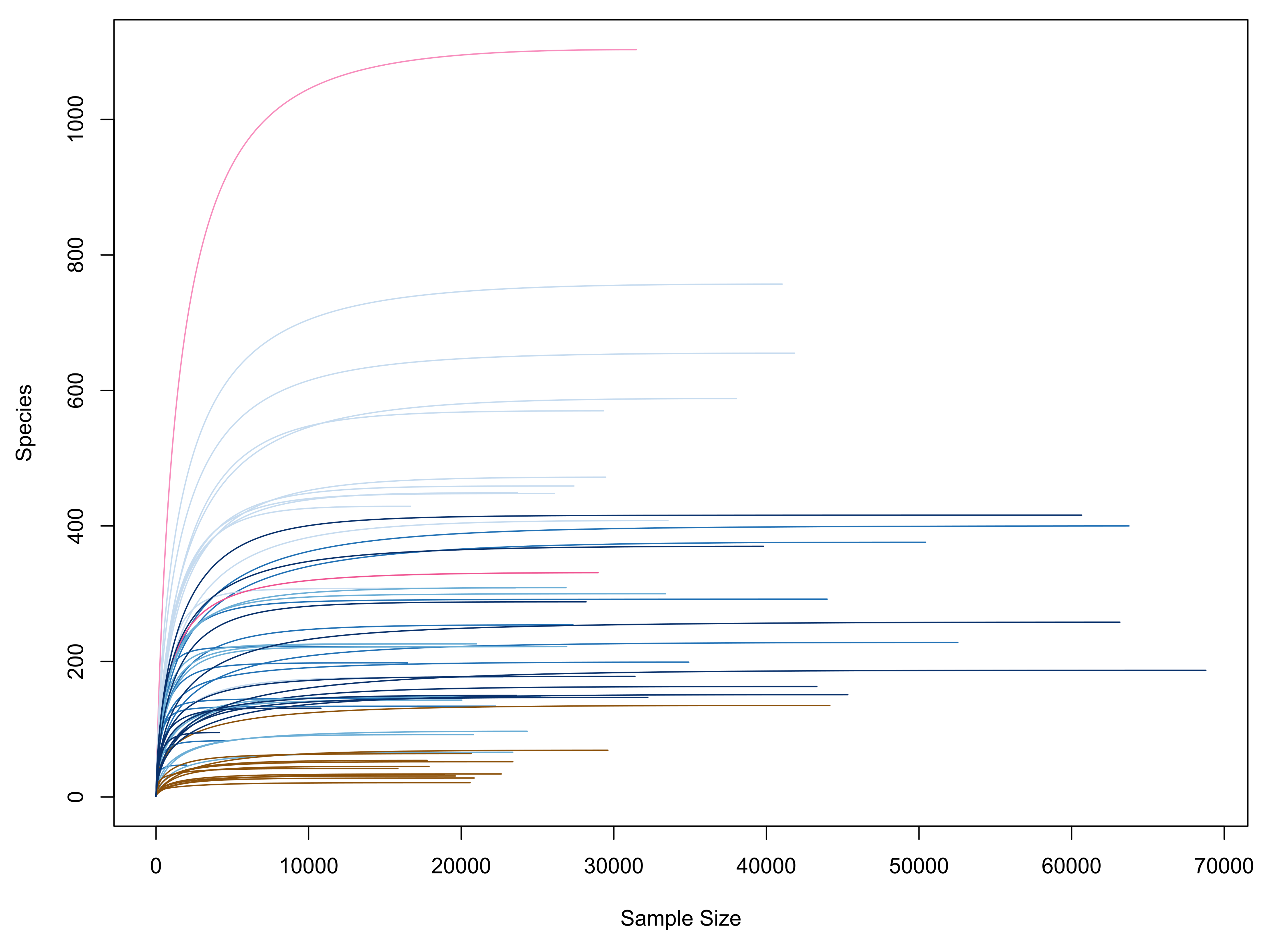

### Supplemental Figure 2

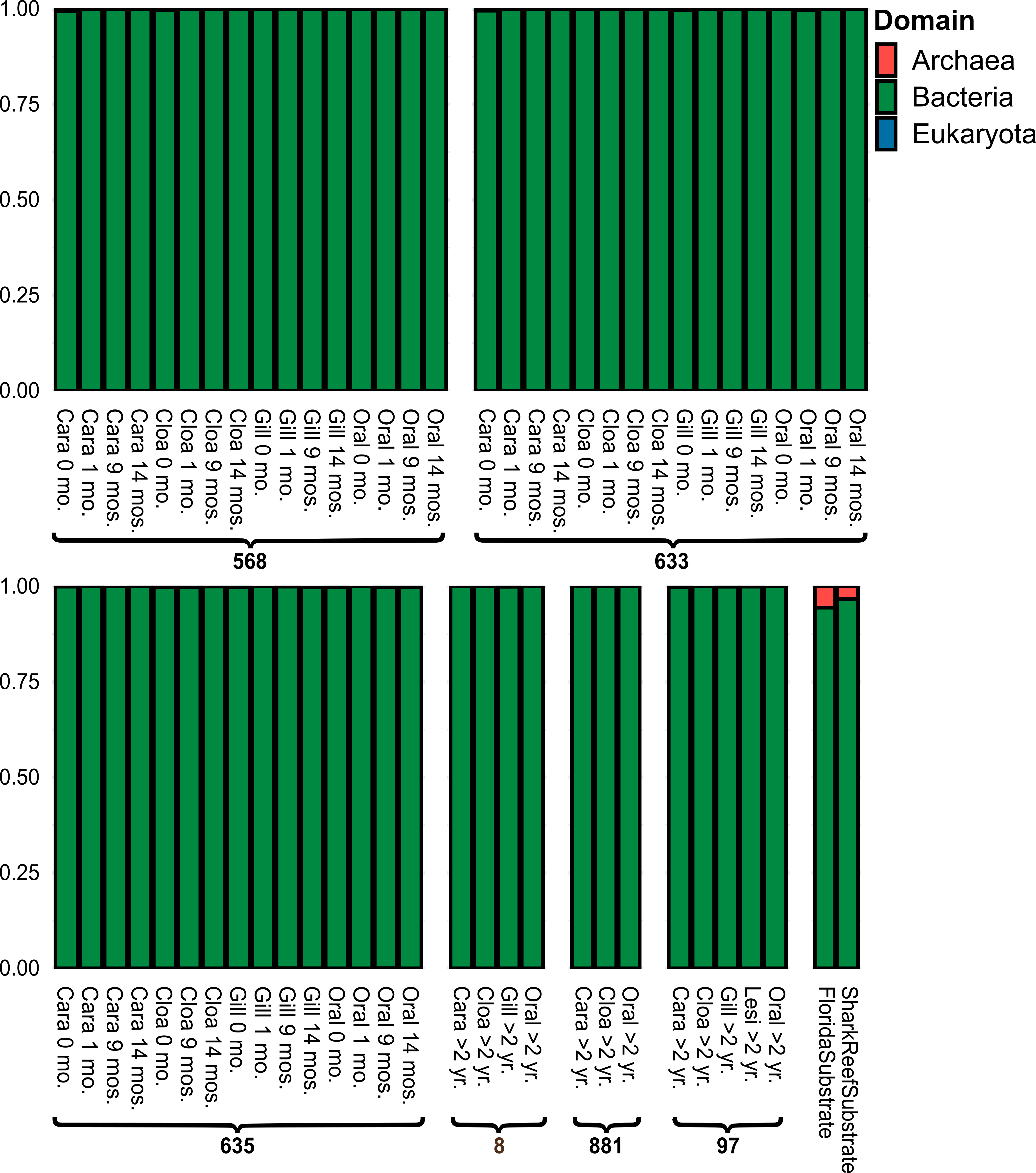

### Supplemental Figure 3

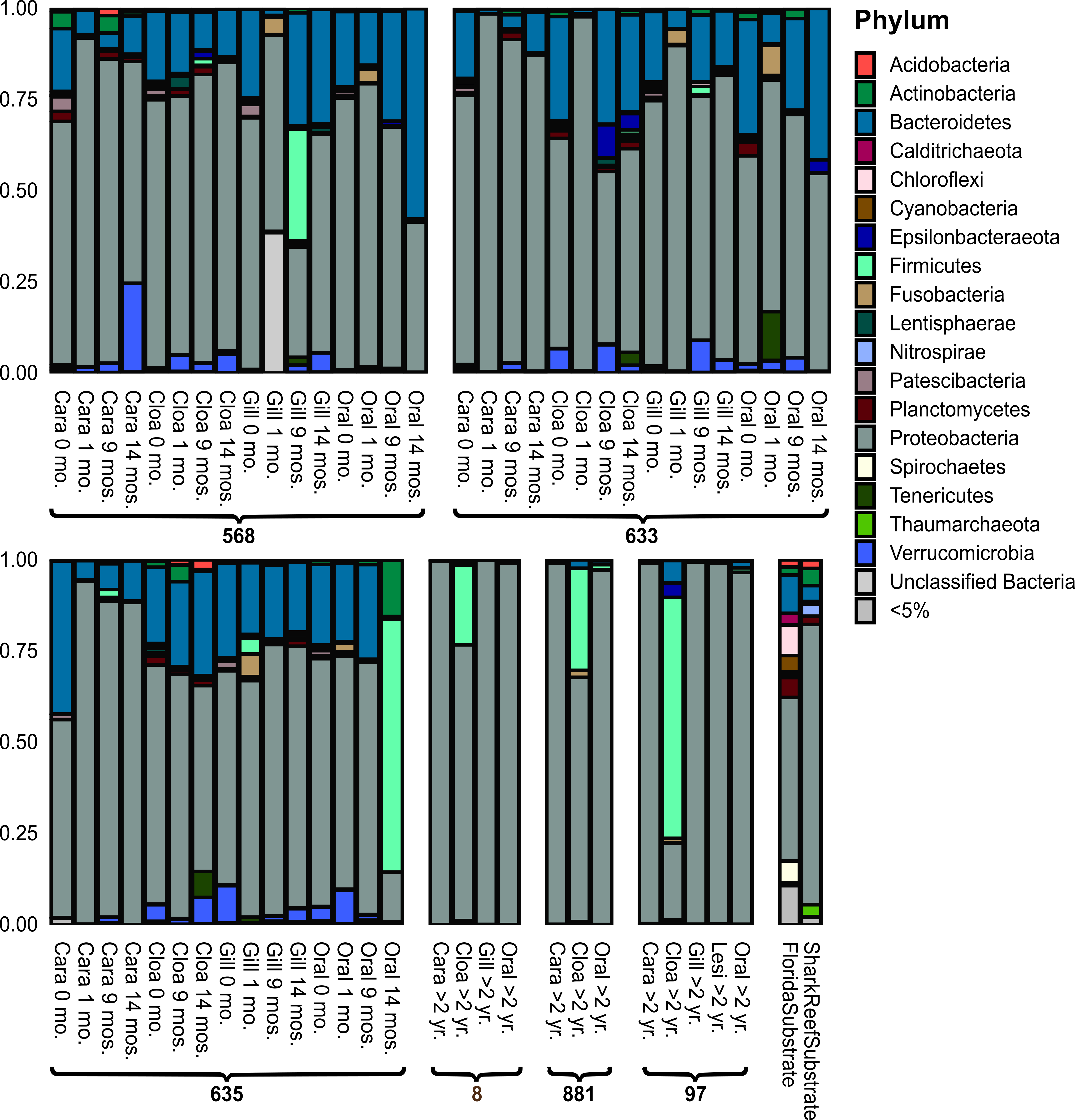

### Supplemental Figure 4

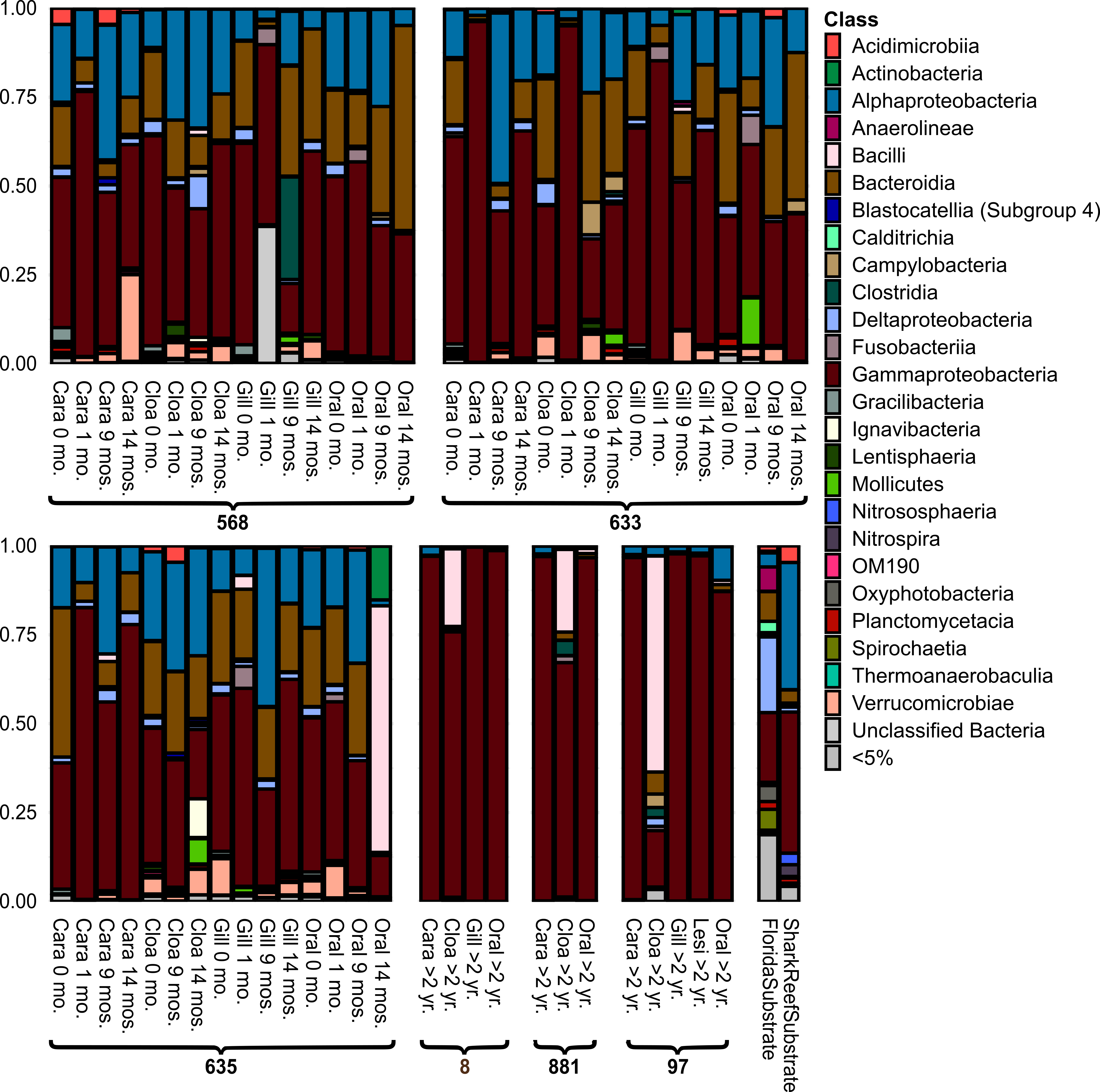

### Supplemental Figure 5

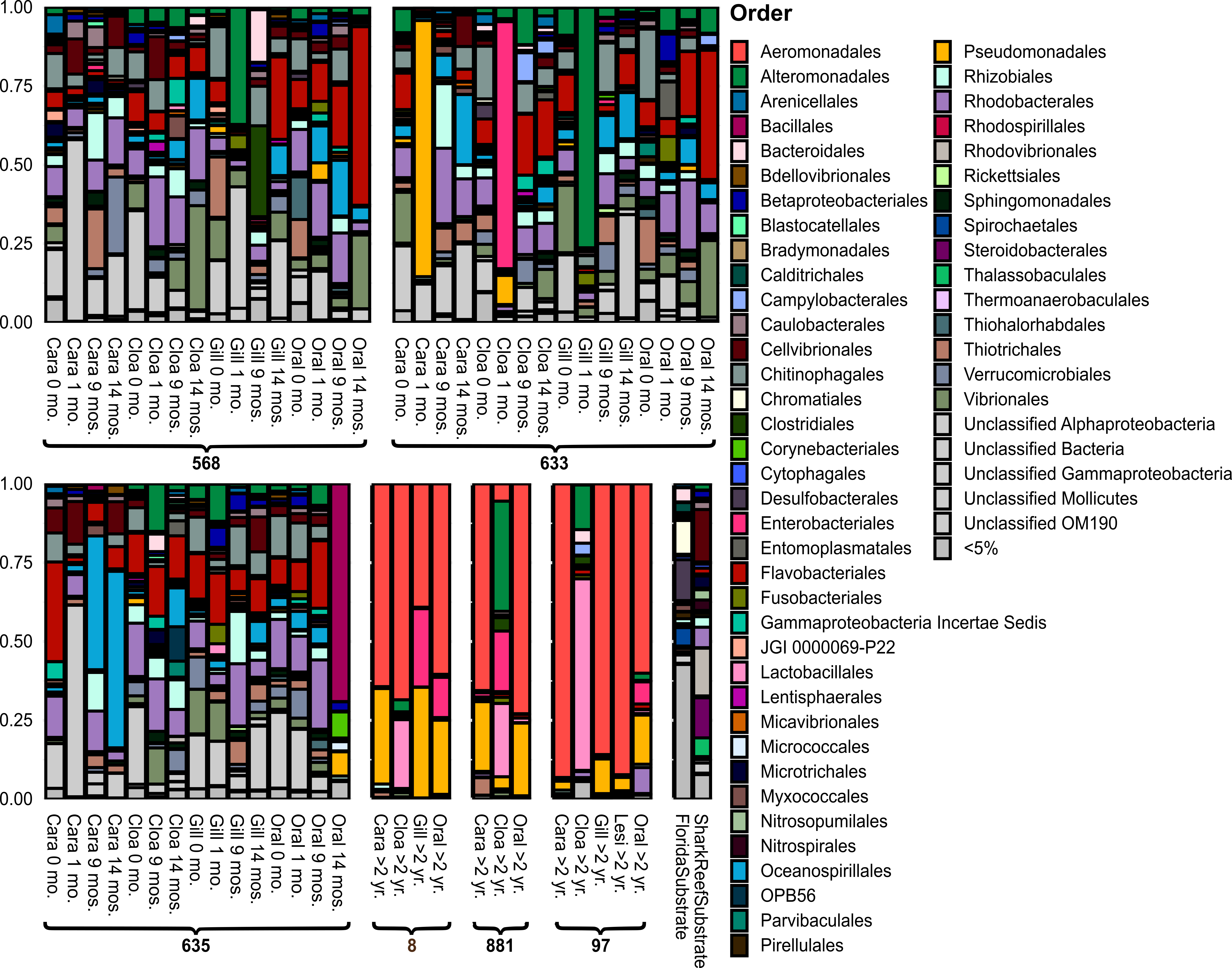

### Supplemental Figure 6

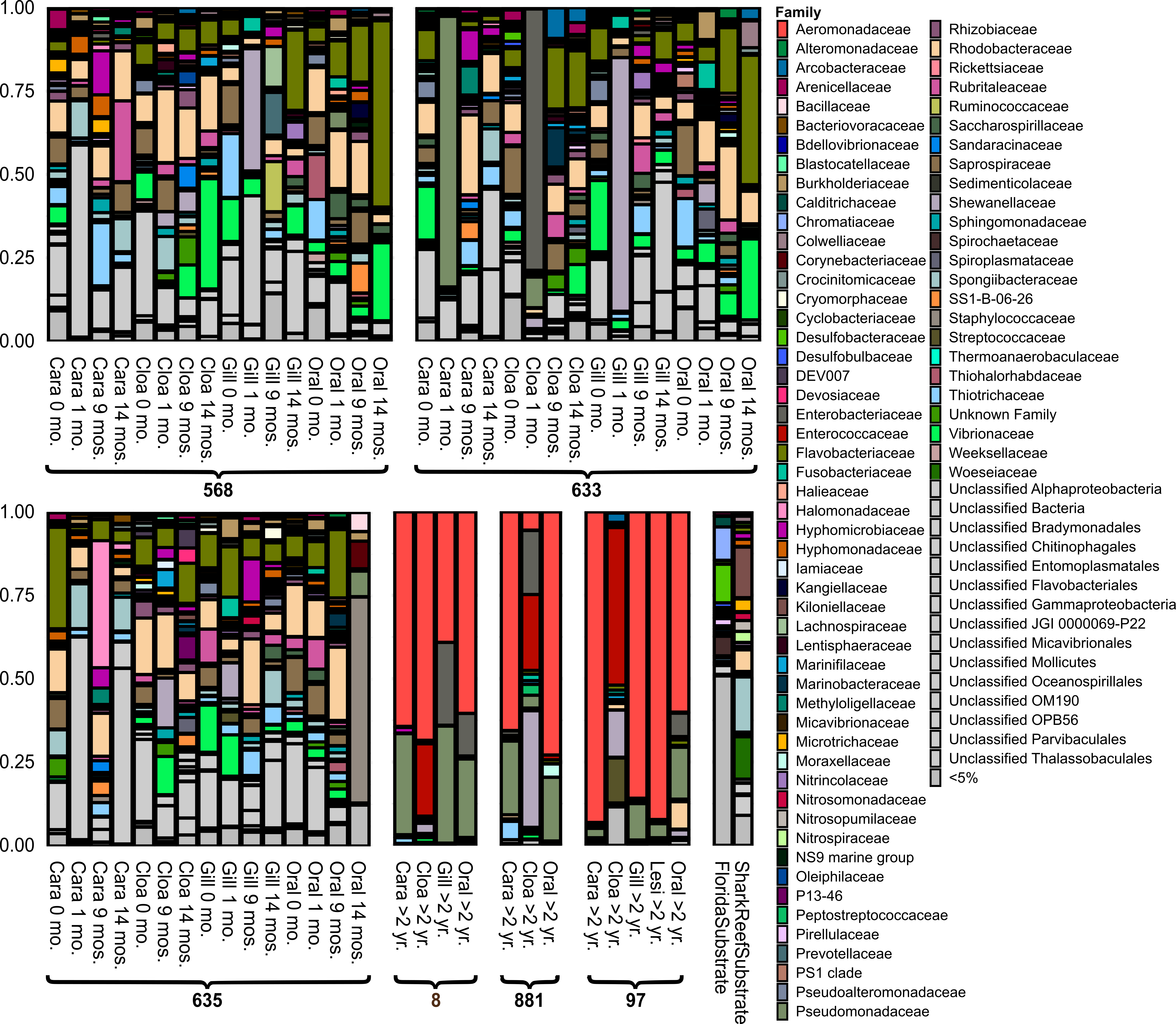

### Supplemental Figure 7

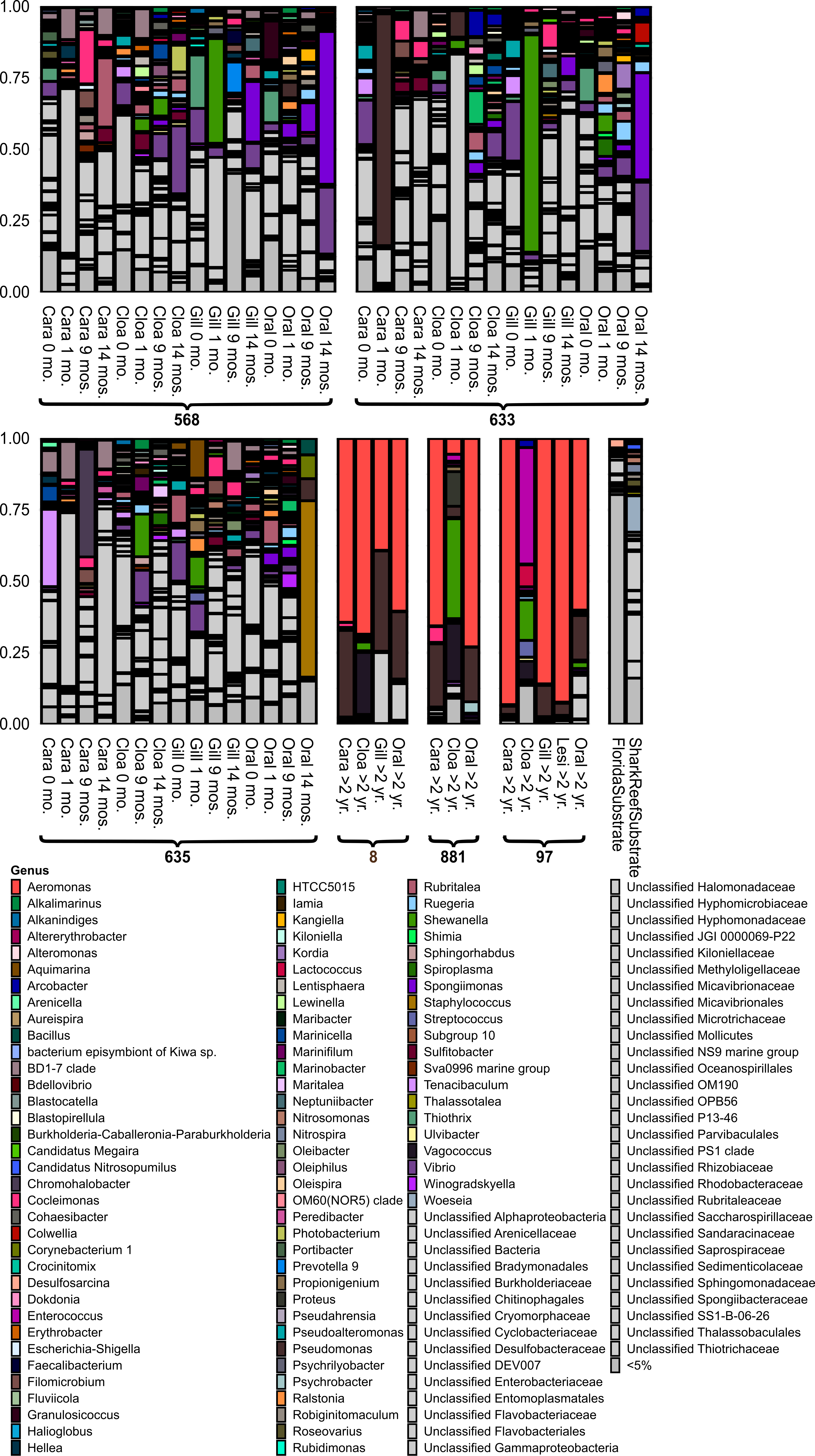

### Supplemental Figure 8

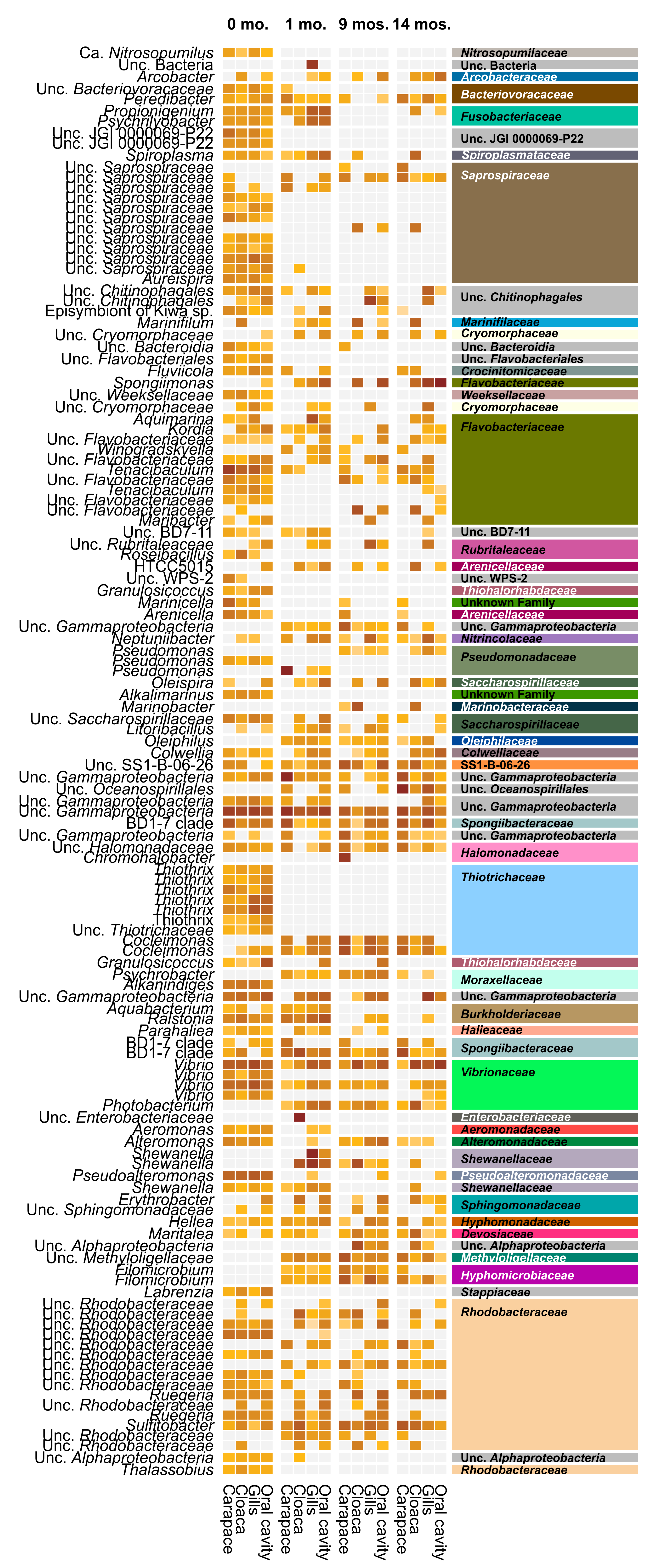
